# Protein and genomic language models chart a vast landscape of antiphage defenses

**DOI:** 10.1101/2025.01.08.631966

**Authors:** Ernest Mordret, Alexandre Hervé, Hugo Vaysset, Tyler Clabby, Florian Tesson, Helena Shomar, Rachel Lavenir, Jean Cury, Aude Bernheim

## Abstract

The bacterial pangenome encodes an immense array of antiphage systems, yet much of their diversity remains uncharted. In this study, we developed language models to predict novel antiphage proteins in two ways: first via fine-tuning ESM2, a protein language model capable of detecting distant homology to known defense proteins, second via a genomic language model with ALBERT architecture which predicts defensive function based on genomic context. We demonstrate that applying these approaches to Actinomycetota - a phylum largely unexplored for antiphage defenses, can accurately predict previously unknown functional defense mechanisms, leading to the discovery and experimental validation of six defense systems with novel antiphage proteins. Analysis of over 30,000 bacterial genomes predicted more than 45,000 uncharacterized protein families potentially involved in antiphage defense, underscoring the vast, untapped diversity of these systems.

---

Bacteria ward off invading phages and other selfish genetic elements through diverse mechanisms^1^. These antiphage defense systems comprise proteins or operons that detect phage incursions and trigger responses that disrupt various stages of the phages’ life cycles^1^. To date, more than 200 antiphage defense systems have been experimentally validated^2^, with dozens characterized in detail, uncovering a remarkable diversity of molecular mechanisms^3–8^.

The rapid expansion in the discovery of defense systems was initially driven by computational approaches based on the observation that defense systems tend to colocalize in bacterial genomes in structures known as “defense islands”^9^. Following the principle of “guilt by association”, protein families frequently found near known antiphage components are hypothesized to play antiphage roles. Experimental validation of these predictions confirmed this hypothesis, and the computational methodologies have since been refined and extended, leading to an explosion in the number and diversity of identified defense systems^10–12^. Beyond defense islands, these systems are often embedded within mobile genetic elements (MGEs), such as prophages and their satellites^13,14^, or integrated into specific loci within genetic structures like integrons^15,16^. Some systems even appear intertwined with others, forming complex “embedded” configurations^17,18^. All of these observations have been harnessed into discovery methods of novel antiphage systems.

Many genes predicted to reside in defense islands remain unexplored, as highlighted by Gao *et al.*^11^ who predicted >7000 protein families enriched in defense islands. Furthermore, even in well-studied model organisms, experimental screens continue to reveal new defense-associated proteins^19^. These findings suggest that the diversity of defense systems could be far greater than currently appreciated. In less-characterized bacterial groups, such as Actinomycetota, the landscape of defense systems remains particularly underexplored. For instance, comparative genomics studies have revealed that these bacteria seem to lack many of the systems commonly found in Pseudomonadota^20^. On the other hand, some defense systems appear to be specific to Actinomycetota, such as lanthipeptide-based systems^21^. Collectively, these observations suggest that the diversity of antiphage defense systems is vast and largely untapped.

Language models, deep learning architectures designed to capture complex relationships in text like data, have recently revolutionized natural language processing^22^. Their key innovation resides in the attention mechanism, which allows the model to capture long-range dependencies and nuanced contextual relationships effectively^23^. In recent years, language models have achieved notable successes in several areas of biology. In particular, protein language models, which leverage natural language processing techniques to analyze protein sequences, have shown remarkable success in predicting protein structure and function^24^. Complementary to this, genomic context-aware models can capture relationships between genes based on their chromosomal neighborhoods, offering new ways to identify functional modules^25–27^. Here, we sought to harness the power of language models to map the landscape of antiphage defense systems.

## Results

### ESM-DefenseFinder predicts putative novel antiphage systems through homology

We first established the extent of known antiphage systems (composed of one or several antiphage proteins). Using the defense system detection softwares DefenseFinder and PadLoc^28,29^, we annotated the defense repertoire of all prokaryotic genomes in RefSeq (n=32,798) (**Supplementary Table 1,2)**. DefenseFinder focuses on experimentally validated systems, while PadLoc also includes systems predicted as antiphage because they are found embedded within known defense systems^17^. Among the 123M proteins in the RefSeq dataset, DefenseFinder identified 511,489 (0.41%) as components of antiphage systems, while PadLoc detected 805,357 (0.65%) (**Figure 1a, Supplementary Table 1,2**). Most proteins annotated by DefenseFinder were also found by PadLoc (83%), while only 52% of PadLoc-detected proteins overlapped with DefenseFinder (**Figure 1a**). Furthermore, Gao et al. published a list of 7,472 protein families predicted to have a defensive role through guilt-by-association methods (presence in genomic contexts enriched in known defense systems). Among them, only 38% were covered by DefenseFinder or PadLoc profiles (**Figure 1b**), highlighting that these tools, which represent our current understanding of antiphage systems, might miss a substantial portion of antiphage proteins diversity.

**Figure 1:**
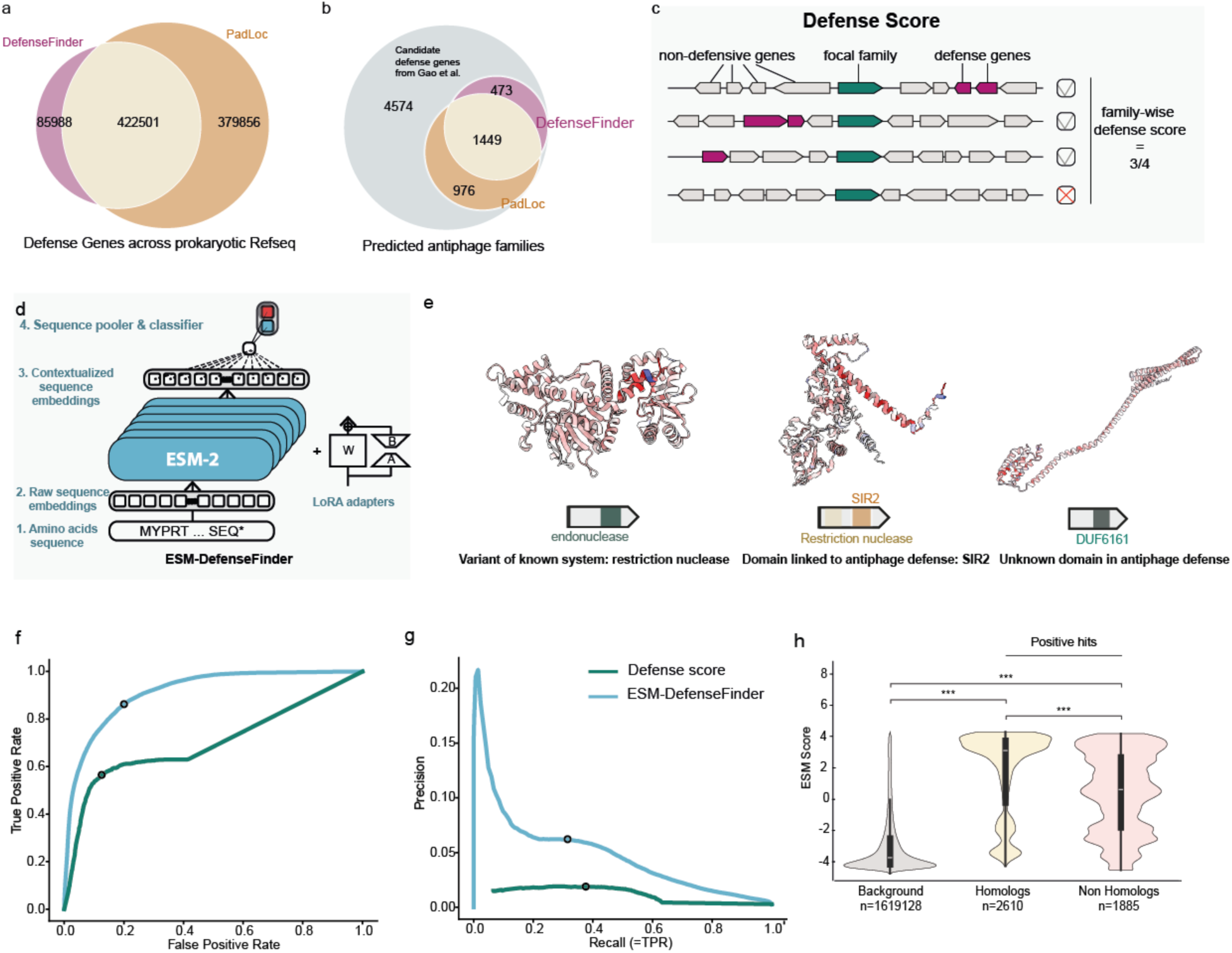
ESM-DefenseFinder predicts novel antiphage systems through homology. **a.** Number of detected defense systems across the prokaryotic RefSeq database, according to DefenseFinder (red) and PadLoc (orange). **b.** Intersection of the Gao et al. candidate proteins with the DefenseFinder and PadLoc profile HMMs. **c.** Family-wise Defense Score calculation, adapted from Doron *et al*^10^. **d**. ESM-DefenseFinder is an ESM2-35M parameters model fine-tuned for defense classification using LoRA adapters. **e** Predicted candidates of ESM-DefenseFinder. Domains were annotated using HHPred. Structures are colored with contribution of each amino acid to ESM-DefenseFinder decision using the layer integrated gradient method^38^. Red means pushing the decision towards defensive **f-g.** Receiver Operator Curve (f) and Precision-Recall curve (g) of ESM-DefenseFinder and Defense score over the test set proteins (see methods). Blue dots represent the performances at the threshold of best informedness (f) and best F1-score (g). At the optimal F1-score threshold for ESM-DefenseFinder (3.27), precision was 9.4%, with a recall of 18% (Methods). Defense score achieved a precision of 2.9% with a recall of 25.3% at optimal F1 threshold (0.18) **h.** Distribution of ESM-DefenseFinder scores for the negative (“Background”) and positives families from the test set, at 80% amino-acid identity. These positive families were further split based on whether they showed marks of homology to positive train set families (“Homologs” and “Non Homologs”). Homology was assessed via a structural search on the predicted structures of the train and test set positive proteins, using FoldSeek. The two-sided Mann-Whitney U test was used to compare the distributions of the populations, with *** indicating a p-value < 0.001.

Many defense systems have been discovered through their genomic association with known systems. This association can be quantified using a family-wise “defense score”, which measures how frequently a given family co-occurs with known defense gene families (the “anchors”) across genomes (**Figure 1c**). This represents the most widely used approach for predicting novel antiphage proteins and was therefore employed as a benchmark for evaluating other methods. We applied this approach to the RefSeq database of prokaryotic complete genomes. We first clustered the 123M proteins in RefSeq into 1,365,092 families (50% sequence identity) and computed the defense score as the number of homologs in the family with known defense proteins in its genomic neighborhood (Methods). Using this method, 13,244 protein families (0.97% of the total number of families) are predicted as antiphage.

Protein language models (pLMs) have demonstrated an impressive ability to capture the molecular function of proteins in classification tasks, such as prediction of enzyme class^30^. We explored whether a fine-tuned pLM could learn to model the diversity of defense proteins based on what was already experimentally validated in order to accurately identify antiphage proteins that remain unknown. To test this, we fine-tuned a small protein language model, ESM2-35M, to classify proteins as antiphage or not (**Figure 1d**, **Supplementary Figure 1.a**). We used protein families clustered at 80% identity identified by DefenseFinder as the positive class (n = 132,567), contrasting them with the remaining families as negatives (including some families with predictions from PadLoc or homology to the Gao et al. proteins). In a naive setup, cross-validation splits positive families randomly, but this approach risks overestimating the performance of the model to identify novel defense proteins as different families from the same defense system could be found in the train and in the test sets. To address this, we split the 137 defense system types identified by DefenseFinder into distinct training, validation, and test sets (**Supplementary Table 3**), and divided their associated proteins accordingly so that all protein families belonging to a given system are found in the same dataset partition. For example, all Lamassu protein families would belong to either training, validation or test (Methods and **Supplementary Figure 1a**). We named the fine-tuned classifier ESM-DefenseFinder and applied it to predict novel antiphage proteins.

Examining candidates predicted by ESM-DefenseFinder **(Figure 1d, Supplementary Table 4)**, we observe that a first group of predicted protein families appear to be variants of known antiphage proteins. For example, many high scoring families are endonucleases, a class of enzymes commonly involved in antiphage defense. A second group of families exhibit partial sequence homology to known defense systems but differ significantly in their characteristics^28^, as exemplified by a *Mucilaginibacter* protein, which bears a SIR2 domain, recently linked to antiphage defense^31^, but associated with a restriction nuclease domain, a previously undescribed combination of domains in antiphage defense. Lastly, a third group contains families for which we could not find any known characteristics of defense systems, for example, a family of proteins found in Pseudomonadota with a DUF6161, with no detectable homology to known defense proteins. Finally, some high-scoring hits correspond to proteins with established non-antiphage functions, such as Ku from the NHEJ repair pathway,which shares homology with csn2, a protein involved in type II-A CRISPR-Cas system^32^, indicating that the method also generates false positives.

We then sought a more quantitative evaluation of ESM-DefenseFinder. ESM-DefenseFinder achieved an AUROC of 73.7% on the test set, a classification performance on par with the one of the defense score on the same set of protein families (78.2%, **Figure 1f**). Since the abundance of defense system types varies drastically, with some like Restriction-Modification or CRISPR-Cas representing large “jackpot” groups of defense proteins as they are the most abundant antiphage systems in the bacterial pangenome, we wondered if the model’s overall performance could be driven by a few well detected and abundant systems but perform poorly on rare ones. To account for this, we studied separately the model’s response over different types of defense systems of the test set (**Supplementary Figure 2a**). The models demonstrated strong performance across all types of defense systems, including rare ones like RADAR. However, certain systems, such as mazEF, were infrequently identified as antiphage, which could reflect that MazEF, a well-known toxin-antitoxin system, is believed to have roles that extend beyond defense^33^. These results suggest that ESM-DefenseFinder effectively assigns high scores to a broad range of antiphage proteins.

Although the high AUROC suggests good overall performance, it does not properly account for false positives due to the strong class imbalance, since there are very few positive labels (proteins annotated as defense systems by DefenseFinder) in the dataset compared to negative ones (>99%). To address this, we evaluated the model using precision-recall curves. The model achieved an average precision of 4.9%, whereas the defense score, by comparison, achieved an average precision of 1.1%. (**Figure 1f-g**). The seemingly low precision may stem from two potential causes. First, these methods can produce false positive predictions: the defense score might capture many proteins associated with mobility (transposases, integrases…), while ESM-DefenseFinder might capture close homologs of antiphage proteins that have a different function. Alternatively, some of the apparent “false positives” could actually be genes involved in antiphage defense that have not yet been characterized as such but are still accounted for as negative examples in our cross-validation setup.

To better understand the nature of ESM-DefenseFinder false positive predictions, we studied high-scoring false positives in *E. coli,* a well characterized bacterial species (**Figure 1g**). ESM-DefenseFinder assigned high scores to YhdJ and MukB (3.418 and 3.5, respectively), proteins with established functions unrelated to defense^34,35^. This could be attributed to their homology with restriction methylases and with LmuB and JetC from the Lamassu and Wadjet systems, respectively. Interestingly, certain proteins, such as an AAA-ATPase from (WP_032269946.1, score = 3.84), are predicted to be antiphage by PadLoc (here PDC-S06), suggesting that these may represent, in fact, true positives. To obtain a more quantitative perspective, we analyzed 2559 genomes of *E. coli*. We computed their core and pan genomes and annotated the proteins using DefenseFinder and PadLoc (**Supplementary Table 5, Supplementary Figure 2b**). Among the 3,496 clusters in the persistent genome (50% identity, present in 90% of genomes), only 15 exceed the high score threshold (0.43%). In contrast, within the *E. coli* pangenome, 3,612 out of 42,876 (8.42%) clusters achieve high scores highlighting that high-scoring proteins mostly belong to the accessory pangenome as expected for antiphage proteins. High-scoring proteins also showed significant enrichment for both DefenseFinder and PadLoc hits (26% compared to 0.4% in the background), as well as hypothetical proteins (26% vs 4%). (**Supplementary Figure 2b**). This suggests that many “false positives” may represent true but uncharacterized defense proteins.

Finally, we seeked to gain a better understanding of what drives ESM-DefenseFinder’s decisions. Based on the observation that high scoring false positives are often found to be homologous to known defense proteins, we hypothesized that the model relies heavily on homology with known antiphage proteins. We suspected that the model would perform better on proteins of the test set that share some sequence homology with the proteins of the train set, as many defense systems share domains—for example, the PARIS defense system contains a AAA_15/21 domain, shared with proteins from the Gabija, AbiL, Retron I B, MADS3-4, PD-T4-4, and Septu systems^36,37^. To test this, we split test set proteins into those with and without detected homologs among training set defenses and compared their score distributions. Homology was assessed using MMseqs2 for sequence similarity (**Supplementary Figure 2.c**) and Foldseek for structural similarity (**Figure 1h)**. In both cases, proteins with antiphage homologs in the training set scored significantly higher than those without. Interestingly, proteins without homologs still scored above the background population (Mann Whitney-U, p<0.001), suggesting that while ESM-DefenseFinder does rely largely on homology, additional features might inform its predictions. Alternatively, the model could be informed by remote homology not detected by our methods.

Overall these findings highlight the potential of ESM-DefenseFinder to uncover novel defense proteins.

### Genomic context-based transformer models identify novel putative antiphage systems in Actinomycetota

ESM-DefenseFinder mainly relies on homology to detect defense proteins and is therefore unlikely to identify proteins whose antiphage mechanisms involve domains or folds not seen during training. To address this limitation, we explored complementary methods that rely on fundamentally different principles. Context-based methods for defense system discovery operate on the principle of “guilt by association,” wherein proteins sharing similar genomic contexts (for example defense island or MGE hotspot) are likely involved in similar biological processes. These methods require a diverse genomic corpus and a finite set of protein families, referred to as the vocabulary. Given the vast diversity of bacterial pangenomes, we focused on a specific bacterial phylum and defined an appropriately sized vocabulary. As most antiphage system studies focus on model organisms like *E. coli*, *B. subtilis*, and *P. aeruginosa*, we selected Actinomycetota—a phylum largely unexplored for its antiphage defenses—as our target.

Actinomycetota, known for their diverse genomes and complex lifestyles, have been heavily mined for secondary metabolites, yet their defense repertoire remains mostly uncharacterized^20^. We compiled a dataset of 10,796 diverse actinobacterial genomes, including many genomes from the genus *Streptomyces* (n=2,880, 26.6%), and clustered their 54 million proteins into families at 50% amino acid identity (n=4,2M, **Supplementary Figure 3a, Supplementary Table 6**). We capped our vocabulary to cover the 2^19^ (∼524k) most common gene families, encompassing 89% of the proteins in the dataset (Methods). The remaining 11% of rare proteins were excluded from the vocabulary. Although these may include potential antiphage proteins, their sparse context is not particularly informative, as 71% of these excluded families are singletons (**Supplementary Figure 3a**).

We first calculated the defense score (see previous paragraph) for all the protein families in the vocabulary to serve as a baseline to compare the performance of the developed predictive methods (Methods). Over a test set comprising 230 positive families coming from 20 defense system types and 36,466 negative families (0.63%), the defense score achieved an AUROC of 80.4% and an average precision of 4.4% (**Figure 2c-d, Supplementary Table 6**). As an additional benchmark, we also implemented a Word2Vec-inspired approach (Methods) which achieved an AUROC of 85.9%. and a similar average precision of 4% (**Figure 2d-e, Supplementary Figure 3b**)^27^.

**Figure 2:**
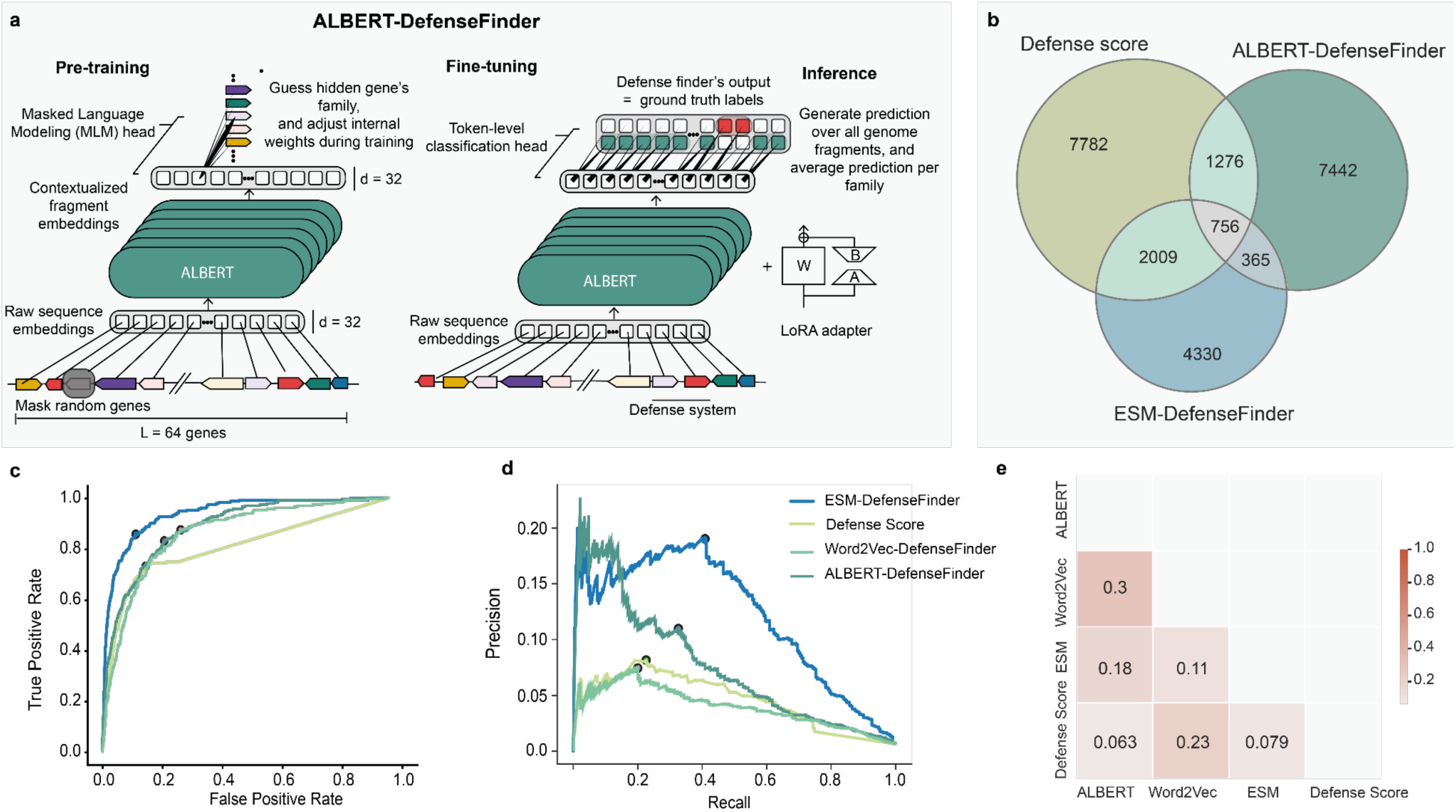
Context-based transformer models identify novel putative antiphage systems in *Streptomyces*. **a.** ALBERT is a Genomic Language Model using protein families as tokens. We first pre-trained it on the task of Masked Language Modeling (MLM), during which masks are applied over randomly picked proteins in the input genome fragment, and the model must “guess” the family of the missing proteins (left). Once pre-trained, the MLM head was swapped for a token classification head, and we fine-tuned the model on the task of token-wise defense protein classification (right). Finally, the model was run in inference mode over all genome fragments from the full Actinomycetota dataset, and its predictions were averaged over genomic positions (ALBERT gene-wise score) and finally aggregated over protein families (90th percentile ALBERT family-wise score). **b.** Venn diagrams comparing the set of protein families from Actinomycetota predicted as antiphage according to the defense score (light green), ALBERT-DefenseFinder (dark green) or ESM-DefenseFinder (Blue). **c-d** Receiver Operator Curve (d) and Precision-Recall curve (e) of the four family-wise scores over the test set protein families of the vocabulary. We evaluated the classifiers ability to retrieve positive (= annotated as antiphage by DefenseFinder) families from a set of negative families. Dots represent the thresholds of best informedness (e, respectively ESM 0.936, ALBERT 0.884, Defense Score, 0.804, Word2Vec 0.859) and precision at best F1-score (ESM 0.123 (2.78), ALBERT 0.077 (2.67), Defense Score, 0.0044 (0.33), Word2Vec 0.040 (2.72)) for each score. **e.** Inter-scores correlation matrix. Spearman correlation coefficient between pairs of scores, over the vocabulary (524k most common actinobacterial gene families at 50% aa identity). ESM-DefenseFinder were first computed over representatives of families at 80% identity, and averaged over families at 50% aa identity.

We then aimed to leverage advanced language model architectures to predict novel antiphage systems using genomic context information. Following a word–protein family analogy^27^, we trained a language model to “read” stretches of bacterial genomes containing 64 contiguous protein families (“tokens”) each. Due to the larger vocabulary in our dataset compared with classical natural language processing tasks (524,288 entries compared to ∼30,000 words in natural language), we employed a modified version of the ALBERT architecture (**Supplementary Figure 3c**, Methods)^39^. Our ALBERT model uses as input: (i) the family (token) assigned to each protein within the genomic fragment, (ii) the relative positions of the proteins within the fragment, and (iii) the orientation of the coding regions (Methods). The ALBERT model, containing 44 million parameters, was first pre-trained on a masked language modeling objective over 6.7 million fragments of 64 genes each (432 million tokens)^40^. We then fine-tuned the transformer, “ALBERT-DefenseFinder”, to classify proteins as antiphage or not.

ALBERT-DefenseFinder achieved an AUROC of 88.4% and an average precision of 7.7%. For comparison, we reran ESM-DefenseFinder on this dataset and obtained slightly higher classification performance (AUROC = 93.6%, average precision = 12.3%) (**Figure 2c-d, Supplementary Table 6**). Despite their similar performance, these methods predict different sets of proteins as antiphage in the test set. Using the thresholds optimized for the best F1-score, only 756 protein families were identified as antiphage by the three models (defense score, ESM-DefenseFinder and ALBERT-DefenseFinder, **Figure 2b**). In contrast, the combined approaches of ALBERT-DefenseFinder and ESM-DefenseFinder predicted an additional 12,137 protein families beyond those identified by the Defense Score alone. Only 463 protein families were predicted as antiphage by all three machine learning methods (ESM, Word2Vec, ALBERT), representing just 1.98% of the 23,344 families predicted positive by at least one method (**Supplementary Figure 4a**). Half of these shared proteins (234) were already annotated as antiphage (DefenseFinder), with greater overlap between context-based methods. Indeed, principal component analysis (PCA) of the four scoring systems revealed that ALBERT-DefenseFinder and Word2Vec-DefenseFinder scores are collinear, while ESM-DefenseFinder scores are orthogonal, highlighting their distinct predictive approaches. Together, the first two principal components explained 76% of the variance (**Supplementary Figure 4b**). These results underscore the complementarity of protein and genomic models in identifying novel defense proteins.

### Experimental validation uncovers six novel actinobacterial antiphage systems

We then sought to evaluate whether language models could uncover antiphage systems radically different from what is currently known. As we had established that ESM-DefenseFinder largely relies on homology with already validated defense systems, we focused on ALBERT-DefenseFinder predictions. To assemble systems from antiphage proteins, we focused on groups of proteins whose co-occurrence in genomic neighborhoods influenced their classification as antiphage by ALBERT-DefenseFinder (Methods). This approach enabled the automatic generation of a list containing over 200 predicted defense systems, which were then manually curated. From this list, ten systems were selected. Interestingly, 9 out of 10 systems had low defense and ESM-DefenseFinder scores. For each system, two evolutionarily distinct homologs were synthesized and cloned under a strong constitutive promoter. These constructs were integrated into the genome of the model strain *Streptomyces albus* to evaluate their antiphage phenotypes (Methods, **Supplementary Table 7**).

To assess their efficacy, we tested the candidate-expressing strains against 12 phages from four genera and compared their performance to a negative control, constructed in the same way but expressing enzymes lacking antiphage activity (Methods and Shomar et al.^21^). Of the 10 predicted systems, two exhibited toxicity in *S. albus* under the tested expression conditions. Among the remaining eight systems, six demonstrated increased resistance to at least one phage with more than 100 fold reduction in plaque forming units (PFU) (**Figure 3a, Supplementary Figure 5**). As *Streptomyces* are mostly soil dwelling bacteria, we named these systems after deities of the soil and earth (Ceres, Geb, Veles, Prithvi, Ukko and Oshun). On average, these systems confer robust protection, such as the Ukko system, which offered a 1000 fold protection against phage Djones4 and Djones7. We noted that some of the protection profiles were similar across phages,This could stem from a limited phage diversity compared to studies in *E. coli* as all phages currently available for S. albus belong to the *Arquatrovirinae* subfamily or to the strong constitutive expression used in this model organism^41^. These results validate ALBERT-DefenseFinder’s ability to predict novel antiphage systems accurately.

**Figure 3.**
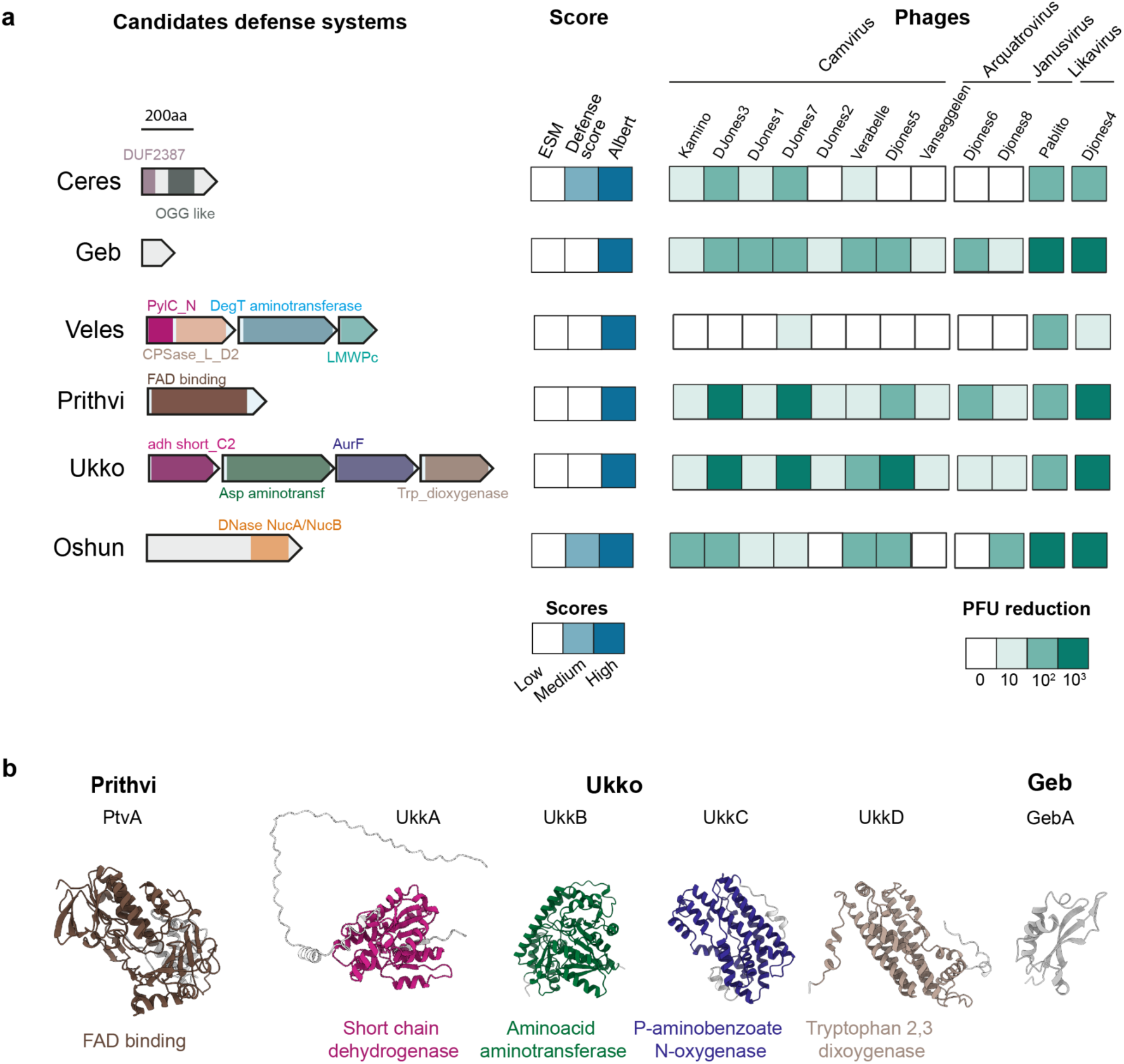
Six antiphage systems identified in *Actinobacterial* genomes display antiphage activity *in vivo*. **a. Candidate defense systems identified in *Streptomyces* genomes using the ALBERT genomic-context transformer model.** Candidate defense systems are predicted from *Streptomyces* genomes using the ALBERT transformer that leverages the genomic context surrounding each gene. Left: Genomic architecture of each system as well as the Pfam domains annotated using HHpred are indicated. Center: Score assigned to each system according to diverse prediction methods (ESM-DefenseFinder: low < 1.13; 1.13 <= medium < 3.27; high >= 3.27, defense score: low < 0.13; 0.13 <= medium < 0.35; high >= 0.286 and ALBERT: low < 0; 0 <= medium < 1; high > 2.67). Right: Defense profile associated with each system when heterologously expressed in *S. albus*. These were established over 3 biological replicates (**Supplementary Figure 5**). **b. Protein models and functional annotations for the Prithvi, Ukko and Geb defense systems.** Models for each component of the Prithvi, Ukko and Geb defense systems are predicted using AlphaFold3 and annotated with the identified Pfam domains using HHpred, as in Figure 3a.

Next, we computationally explored the potential mechanisms underlying the observed antiphage activities. Each protein from the validated systems was modeled using AlphaFold3^42^ and functionally annotated with HHpred^43,44^ and Pfam^45^. Notably, most annotated Pfam domains had not been previously described in known defense systems (**Figure 3a-b**). For instance, the Prithvi system is annotated as a FAD/NAD-dependent aromatic cycle halogenase (PF01494, PF04820, PF05834, PF01266), suggesting that FAD or NAD could act as a cofactor of a novel enzyme involved in antiphage defense (**Figure 3b**). The Ukko system comprises four proteins, all predicted to have metabolic activity. UkkA is annotated as a 2,3-dioxygenase that cleaves the pyrrole ring of tryptophan (PF03301, PF01231, PF08933). UkkB is an aminotransferase (PF12897), UkkC is homologous to AurF 4-aminobenzoate N-oxygenase (PF11583) and UkkD is a NAD(P)-H-dependent enzyme (PF13561). We hypothesize that these proteins catalyze sequential reactions in a biosynthetic pathway that uses tryptophan as a potential precursor to inhibit phage infection (**Figure 3b**). One intriguing system, Geb, contains a single 150-amino-acid protein with no annotated Pfam domains. The protein’s well-defined globular predicted fold and specific histidine and aspartate residues suggest an enzymatic function, though its exact mechanism remains elusive (**Figure 3b**). We then assessed the prevalence of these systems across the >31,000 bacterial genomes of the prokaryotic RefSeq dataset. All systems were rare (less than 200 homologs across RefSeq) and most restricted to *Actinomycetota* (except for Prithvi and Veles, **Supplementary Figure 6**). Overall, these results demonstrate that genomic language models can identify novel antiphage systems with putative mechanisms of action that could differ significantly from those previously described.

### The landscape of predicted antiphage proteins reveals a vast uncharted molecular diversity

We then sought to estimate the global diversity of antiphage proteins. To begin, we assessed this diversity within the genomes of Actinomycetota, where all our methods had been applied. While DefenseFinder identifies 3,171 protein families, combining the predictions of ALBERT-DefenseFinder and ESM-DefenseFinder (union of the two sets of predicted antiphage proteins) suggests that 12,114 novel protein families could be involved in antiphage defense in this phylum and are yet to be experimentally validated. This represents a 4-fold increase compared to what is currently known. More specifically, ALBERT-DefenseFinder accounts for 72% (n = 8,841) of the predicted novelty. This suggests that most of the diversity of antiphage proteins probably remains to be characterized in Actinomycetota.

As an important diversity of antiphage proteins probably also remains to be characterized in other bacterial phyla, we then wanted to extrapolate these predictions to the comprehensive RefSeq database of complete prokaryotic genomes. We thus applied the defense score and ESM-DefenseFinder on this database. However, extending ALBERT-DefenseFinder to the full RefSeq database was not achievable due to inherent limitations of the approach: the vocabulary size of ALBERT-DefenseFinder scales with the number of protein families of the database (n=524,288 on the instance of ALBERT-DefenseFinder). On RefSeq, the vocabulary size of such a model would reach beyond 1,5 million entries. As a consequence it was not possible to train an instance of ALBERT-DefenseFinder on the full RefSeq database of complete prokaryotic genomes.

We first aimed to visualize and interpret this diversity by mapping the predictions onto a protein universe, represented by families of proteins clustered at 50% identity. We selected ESM-DefenseFinder embeddings, projected them into two dimensions, and annotated the resulting UMAP with known defense systems, predictions by ESM-DefenseFinder, or defense score. This UMAP provides an interactive tool to explore the space of predicted antiphage proteins (**Supplementary Figure 7**, mdmparis.github.io/antiphage-landscape/). It reveals clustering of known antiphage protein families (annotated by DefenseFinder) and highlights numerous points predicted by ESM-DefenseFinder and Defense Score that extend beyond well-characterized defense systems, indicating a vast and largely unexplored molecular diversity. Zooming into specific families reveals hallmark features of true antiphage systems. For example, one point reveals a potential two-gene system in the *Vibrio* genus, consisting of a homolog of the HipA toxin and an unannotated gene, both observed in a defense context (**Figure 4a**). Another cluster highlights homologs in *Cyanobacteria* that, while not associated with a defense context, exhibit a genetic architecture reminiscent of NLRs^46^, characterized by a C-terminal repeat domain, a central STAND domain, and an N-terminal region containing an unknown domain (DUF5112) (**Figure 4a**). These results visually encapsulate the opportunity to uncover novel defense systems by combining diverse predictive approaches and provide the community with an interactive way to explore the space of predicted antiphage proteins.

**Figure 4.**
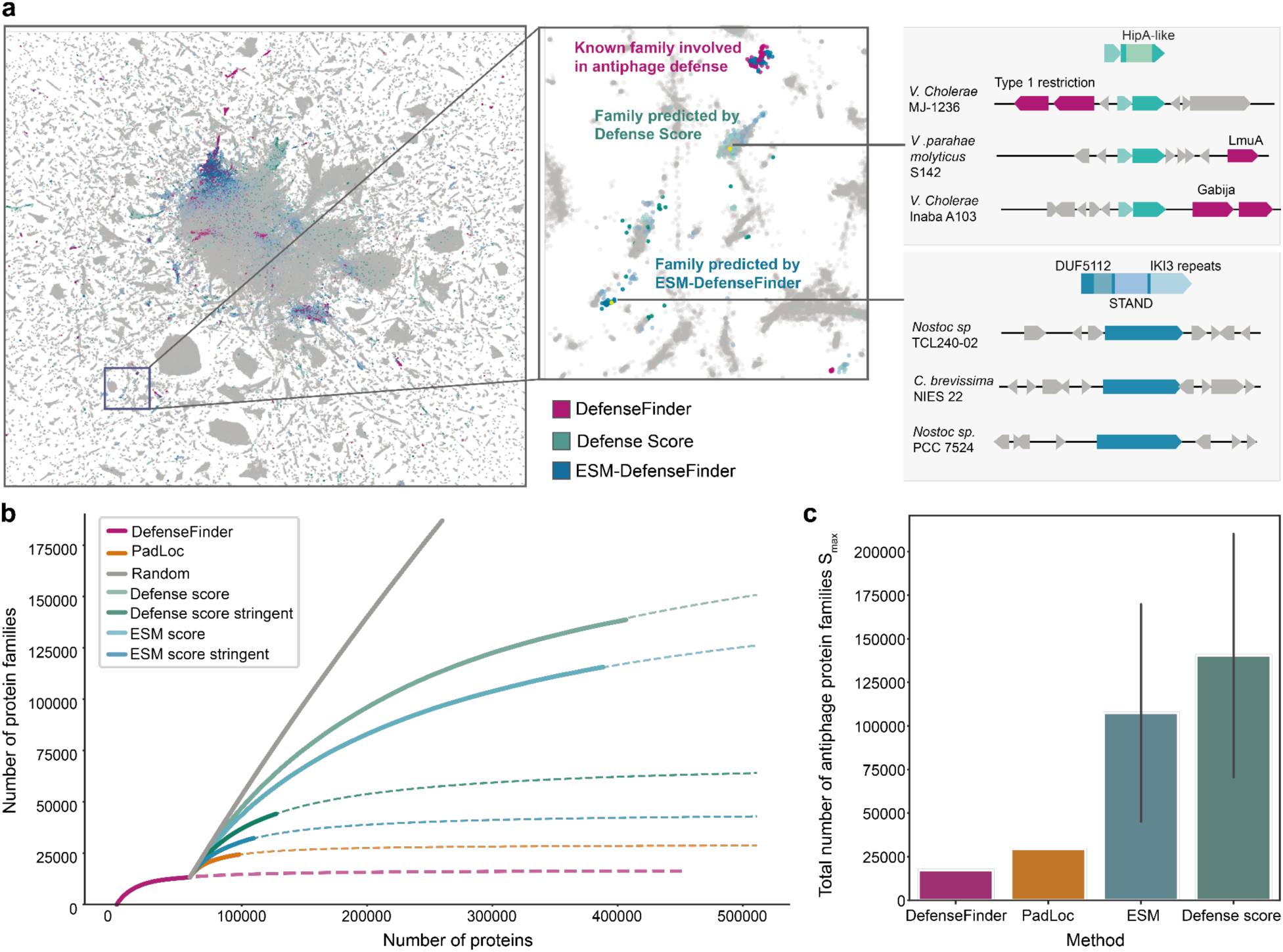
The vast uncharted diversity of predicted antiphage proteins. a. Annotated UMAP of the latent space of ESM-DefenseFinder Background map is a projection of ESM-DefenseFinder embeddings of families clustered at 50% identity, 50% coverage with more than 10 members (n=5,954,941, full map in **Supplementary Figure 7**). Pink: families with a member annotated by DefenseFinder. Blue: families predicted as antiphage by ESM-DefenseFinder (dark blue, score >99th percentile, light blue score >95th percentile). Green: families predicted as antiphage by Defense Score (dark green, score >99th percentile, light green score >95th percentile)**. b. Rarefaction curves of the number of protein families involved in antiphage defense according to diverse prediction methods.** Rarefaction curves assess the number of discovered protein families as new candidate antiphage homologs are discovered. The rarefaction curves are first provided for the experimentally validated antiphage proteins (DefenseFinder, pink). Then the set of proteins not identified by DefenseFinder and detected by PadLoc (orange) or predicted as antiphage by ESM-DefenseFinder (stringent prediction: dark blue, loose prediction: light blue), defense score (stringent prediction: dark green, loose prediction: light green) and for a random baseline (gray) are also provided. For each method, a modeled rarefaction curve obtained by fitting a Michaelis-Menten on the model on each empirical curve is drawn in dashed lines. **c. Estimated total number of protein families involved in antiphage defense per prediction method.** The estimated total number of protein families involved in antiphage defense by each method corresponds to the asymptotic value (Smax) of its corresponding rarefaction curve. For each predictive method, the whiskers correspond to the range between the stringent (score higher than the 99th percentile of the overall distribution) and the loose (score higher than the 95th percentile of the overall distribution) prediction.

We then sought to estimate quantitatively how diverse the set of proteins yet to be described in antiphage immunity is compared with the set of already known antiphage proteins. To do so, we built rarefaction curves from protein families to assess how many novel families are discovered as additional antiphage proteins are identified. Known antiphage protein families account for only 0.97% of all families (n=13,244). Strikingly, the rarefaction curve for these protein families reaches an asymptotic value, suggesting that most known antiphage proteins belong to a closed set of families, with limited additional diversity to uncover (**Figure 4b-c**).

For proteins not identified by DefenseFinder, we used PadLoc, Defense Score and ESM-DefenseFinder to predict antiphage proteins. Rarefaction curves for these methods all reveal additional diversity beyond DefenseFinder predictions (**Figure 4b, Supplementary Figure 8**). To estimate the overall diversity of antiphage proteins, we used the asymptotic values (Smax) of the rarefaction curves. The defense score method consistently identifies more families (Smax = 70,623 for stringent detection, 210,245 for loose detection) compared to ESM-DefenseFinder (Smax = 45,065 for stringent detection, 169,832 for loose detection) (**Figure 4c**). We therefore estimate that between ∼45,000 (ESM-DefenseFinder, lower bound) and ∼216,000 protein families (defense score, upper bound), representing at least 3% of the total protein families analyzed, could be involved in antiphage immunity. Given that DefenseFinder identifies ∼13,000 antiphage protein families, these results suggest that the majority of antiphage protein diversity remains untapped.

## Discussion

Our study demonstrates the power of integrating protein and genomic language models to uncover novel antiphage systems. ESM-DefenseFinder excels at identifying distant homologs of known systems, while ALBERT-DefenseFinder provides complementary insights by leveraging genomic neighborhoods to identify systems with no detectable homology to known defenses and low defense scores. Together, our approaches identified ∼13,000 of putative antiphage protein families within Actinomycetota (**Supplementary Table 6**) and led to the validation of six novel systems. When applied to the bacterial pangenome, these methods uncover an unexpectedly diverse repertoire of putative antiphage proteins, which we estimate to encompass between 45,000 and 216,000 distinct protein families.

Each of the methods applied in this study has specific strengths and limitations. Protein language model ESM-DefenseFinder performs well in identifying proteins with distant homology to known defense systems. However, its reliance on homology limits its ability to discover systems with entirely novel mechanisms. Conversely, the genomic language model ALBERT-DefenseFinder identifies novel systems, even when no homology exists, but is computationally demanding. The complementarity of these approaches is evident in their ability to predict distinct sets of antiphage genes. This synergy highlights the value of combining diverse computational frameworks to achieve a more comprehensive understanding of bacterial defense. A common limitation of all approaches evaluated is the precision which remains very low. This hinders the speed of discovery of novel systems which still requires manual curation. Importantly, our work shows that there is room for improvement for these methods by using more domain-knowledge in the training procedure. We also established a quantitative framework and a set of metrics that can serve as a benchmark for comparing future computational methods.

The discovery of novel defense systems in *Streptomyces* underscores the unique potential of studying non-model organisms. Notably, these systems exhibit associations with metabolic activities, reflecting the rich biosynthetic capabilities of this genus. From an evolutionary perspective, we hypothesize that these systems emerged through exaptation, where components of the metabolic machinery were co-opted for immune functions. Most studies on antiphage systems have focused on a limited set of model organisms, such as *E. coli* and *B. subtilis*. While these organisms have been foundational, they represent only a fraction of the diversity present in nature. By targeting underexplored taxa like *Streptomyces*, we uncover systems that expand the boundaries of known immunity and provide a richer understanding of the strategies bacteria use to combat phages.

This study provides the research community with a list of ∼216,000 candidate antiphage protein families (**Supplementary Table 4 and 6**), an interactive UMAP and predictive models to evaluate whether a protein is potentially defensive. These resources, already available as scripts, could in the future be integrated in web services such as DefenseFinder, enabling streamlined annotation and discovery. These predictions remain hindered by low precisions of the methods and should be manually inspected before experimental validation, which can be performed using existing frameworks such as IMG or Gaia^25,26^. By offering both datasets and computational tools, we aim to accelerate the experimental validation of antiphage systems and facilitate their functional characterization.

The remarkable diversity of antiphage proteins uncovered in this study underscores the complexity of bacterial immunity. This diversity likely reflects the dynamic co-evolution between bacteria and phages, driven by selective pressures to innovate and adapt. Understanding the evolutionary forces underlying this diversity remains a major challenge. Our findings offer a roadmap for future exploration of antiphage systems across the microbial world.

## Materials and Methods

### RefSeq dataset annotation

We downloaded the prokaryotic complete genomes of RefSeq^47^ in May 2023 (n=32,798) with its annotations, and scanned their genomes with DefenseFinder 1.3.0^28^, and PadLoc v2.0.0^29^ to annotate known defense systems. We clustered the full proteome at the increasing granularities of 99%, 95%, 80%, 50% and 30% using MMSEQS2’s linclust command (version 14.7e284) ^48^. For brevity, we will use the notation fam <GRANULARITY> to denote families of proteins at a given granularity, for example fam80 for a family of proteins at 80% granularity.

### Actinomyces Dataset generation / description

We downloaded all n=49,531 actinobacterial genomes from the Secondary Metabolism Collaboratory database (https://smc.jgi.doe.gov/), and complemented them with 11,358 strains from the Shen lab collection (https://npdc.rc.ufl.edu/genomes/). We eliminated redundancy using MASH^49^, using a threshold of average nucleotide identity (ANI) < 0.99, and keeping the genomes with the longest median contigs among the redundant pairs. After eliminating poor contiguity genomes (less than 80% of bp in contigs of size > 50kb, or more than 10^-5^ N/bp among the contigs with size > 50kb), we re-annotated the 10,796 remaining genomes with Prokka^50^. Proteins from the selected genomes were clustered using mmseqs-cluster^48^ at 80%, 50% and 30% identity and 80% coverage using a cascaded clustering strategy.

### DefenseFinder and PadLoc comparison

DefenseFinder v1.3.0 with models v1.3.0 and Padloc v2.0.0 with models v2.0.0 were run on all the RefSeq prokaryotic databases and all hits were compared. All the candidate cluster gene sequences from Gao et al.^11^, were retrieved in the supplementary data. These sequences were scanned with all profile HMM from DefenseFinder and PadLoc using hmmsearch v3.2.2^51^ at the appropriate detection threshold (GA score and coverage for DefenseFinder and E-value and coverage for PadLoc).

### Defense System types split

We randomly split the 137 defense system types of DefenseFinder into three sets: train (94 systems), validation (23 systems) and test (20 systems). We consistently used the same split of defense system types across all analyses.

### ESM-DefenseFinder classifier fine tuning

We created a classification dataset by selecting positive and negative fam80s, each represented by a sequence from a randomly chosen family member. A family was considered positive if at least one member was annotated as defensive by DefenseFinder, and negative otherwise. The dataset was divided into training, validation, and test sets. For each set, we selected positive sequences from families matched by the associated system types. To decrease the prevalence of common defense systems and increase diversity, we sampled with replacement 100,000 training, 5,000 validation, and 5,000 test positive sequences with a probability 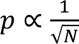, where N is the number of fam80s associated with a given defense gene HMM profile. We split the negative families into three disjoint sets of fam30s for training, validation, and test, and drew 100,000 training, 5,000 validation, and 5,000 test negative sequences to create balanced datasets with an equal number of positive and negative labels.

We fine-tuned an ESM-2 35M parameter model using HuggingFace’s implementation of the LoRA^52,53^ technique for sequence classification. We performed hyperparameter optimization using grid-search to maximize the AUROC over the validation dataset. The hyperparameters included:

- Learning rate: [1e-4, 3e-5, 1e-5]
- LoRA rank: [64, 32, 16]
- LoRA factor: [0.5, 1, 2], such that LoRA_α_ = LoRA rank * LoRA factor

Optimal performance was achieved after training over a single epoch with a learning rate of 1e-5, LoRA rank of 64, and LoRA_α_ of 128. We then trained a model with these hyperparameters on the combined training and validation datasets. We evaluated the model on a variant of the test set. We took all positive proteins from the test set (N positive = 57111) and randomly sampled negative proteins such that the proportion of defense proteins with the test set matches that of the entire database (0.45%, N negative ∼ 12M). We then remove redundancy at 80% of amino-acid identity to prevent over-sampled clades to bias the results. The precision-recall and ROC curves along with their associated statistics were computed on this latter set.

### ESM-DefenseFinder homology scoring analysis

We searched for homology between proteins of the positive test set (i.e. proteins from the test set labelled as defensive) and those of the positive train set, by sequence and structure. We used MMseqs easy-search^48^ (version: 16.747c6) with sensitivity 7 and e-value threshold 1e-3 to search for homologs by sequence. To search for homology by structure we folded the positive proteins of the train and test sets with ESM-3^54^ and searched with Foldseek^55^ easy-search (version: 9.427df8a), using a sensitivity of 13 and an e-value threshold 1e-3. For the two approaches the positive test set was split between those with homologs in the train set and those without. The distributions of scores from homologs, not homologs, and the population were then plotted. The population consists of all the representatives at 80% identity and 80% coverage of the RefSeq database. Significance tests between the distributions were computed with a Mann-Whitney U test, two-sided alternative, for each pair of distributions among: Homologs, Non homologs, and the population.

### Vocabulary construction

We constructed our vocabulary around the most frequent fam50s in the Actinomyces dataset. Additionally, we included non coding genes, such as 64 tRNA tokens (one per anticodon) and 6 rRNA classes, and the special tokens [MASK], [PAD] and [UNK] to represent masked genes, padded genes and unknown (rare) genes. We capped vocabulary size at 2^19^ = 524,288 entries for computational efficiency.

### Unified Actinomyces defense genes classification dataset

In order to compare different methods on equal grounds, we devised a shared dataset to assess their ability to retrieve unseen defensive gene families from a background of non defensive genes. We split the fam50 tokens of the Actinomyces vocabulary into positive and negative families. A family was considered positive if more than 10% of its constitutive fam80s contained at least one defensive member. A family was considered negative if none of its members were matched by DefenseFinder, therefore excluding families with a very low but positive fraction of defensive genes. Then, positive fam50s were split into train, validation and test following the split of defense types described above. All negative families were then split into train, validation and test in a way that guaranteed that the positive family ratio remained constant across folds.

### Defense-score calculation

Defense-scores were calculated using a procedure adapted from Doron *et al.*^10^. We used a granularity of 95% amino-acid identity for C1 clusters, and of 50% amino-acid identity for C2 clusters, so that we could eventually compute a defense-score for each fam50 member of the vocabulary. It is computed as a two-step procedure. First, for each fam95, we compute the fraction of its members that have at least one defense gene in its genomic neighborhood (+/- 10 genes around the focal gene), as detected by DefenseFinder, which we define as the C1 defense score. Then, we average the C1 scores over all the fam95 members of a fam50 to get the C2 defense-score, or fam50 defense score. In order to evaluate this method’s ability to find new defense genes on par with the other methods, we actually computed a partial defense score for each fam50, in which defense proteins that were associated with defense system types of the test set were removed from the list of defense genes before the calculation.

### Word2vec pre-training

We trained Word2Vec embeddings of shapes {vocab_size, 32} and {vocab_size, 300} using a custom implementation of SkipGram^56^. SkipGram generates contextual embeddings by optimizing, over every focal gene *w_I_*, the following Negative Sampling Loss over pairs of neighboring (left hand side) and randomly chosen (right hand side) genes:

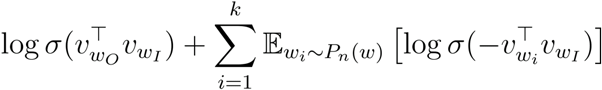

Where:

- *v_wO_* is the embedding of the context gene *w_O_*.
- *v_wI_* is the embedding of the focal gene *w_I_*
- *v_wi_* is the embedding of a randomly sampled gene *w_i_*.
- *σ* is the sigmoid function.
- *k* is the number of negative samples (we chose k=16).
- *P_n_*(*w*) is the noise distribution for sampling negative genes.

Here, we chose *P_n_*(*w*) to follow the unigram distribution U (w) raised to the 3/4rd power, following the original implementation. To enhance the efficiency of the Word2Vec training process, we employed a variant of the Skip-Gram model where the sampling of context pairs and random pairs of gene families was performed in advance, and optimized the loss over large batches (d=16,384) of “neighboring” and “random” pairs of genes.

### Word2vec-DefenseFinder classifiers

Following the approach of Miller *et al.*^27^, we fine tuned a series of Multi Layer Perceptrons (MLPs) to classify all fam50 in the unified Actinomyces defense genes classification dataset. We used a fixed architecture consisting of an embedding layer (output of Skip-Gram), followed by 4 layers with ReLU activation with dimensions [256, 128, 64, 2]. Dropout (p=0.2) was used during training for regularization. The embedding layer was initialized with the pre-trained Word2Vec weights, in dimension 32 or 300, and remained frozen during training. We explored several strategies of class re-weighting (because of class imbalance), in which the weight assigned in the cross entropy loss to each positive example was either untouched (‘flat’), re-scaled by a factor *f*^-1^, where f is the fraction of positive examples in the dataset (‘inverse frequency’), or by a factor *f*^-1/2^ (“square root inverse frequency”). We performed hyperparameter optimization using grid-search to maximize the AUROC over the validation dataset. The hyperparameters included:

- Pre-trained embedding dimensions: [32, 300]
- Learning rate: [1e-4, 1e-5, 1e-6]
- Strategy = [‘flat’, ‘inverse frequency’, ‘square root inverse frequency’]

We used a weight decay of 0, a batch size of 16,384, and an ADAM optimizer. Finally, we selected the best hyperparameters (learning rate = 1e-4, strategy = ‘inverse frequency’, pre-trained embedding = 32), and re-trained a final classifier using the concatenation of train and validation examples. We transformed the logits associated with the positive class, taking first the softmax to derive class probabilities, and then re-transformed them into centered logits using the log odds transformation 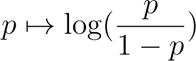 to generate the final Word2Vec fam50 score.

### ALBERT-DefenseFinder architecture

#### Geometric attention

We based our ALBERT model on the HuggingFace implementation, and developed a variant of relative key attention^57^ to more closely capture the geometry of the DNA strand. In the original implementation of the BERT transformer, the output of each attention layer is computed as

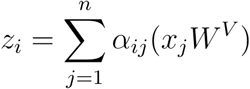

Where α*_ij_*, the attention weights, between tokens i and j are computed as:

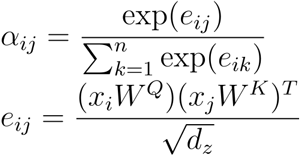

In relative key attention, the last equation is modified to include a column vector 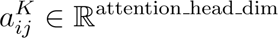, whose values are stored in a learnable lookup table accessed using the clipped difference between the i and j indices.

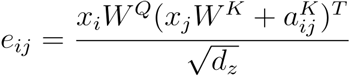

Here, we modified the computation of the 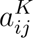 terms, so that it becomes a function of the local coordinates of genes i and j, taking their start and end coordinates as inputs. First, we compute four relative distances for each pair (See Supplementary Figure 3):

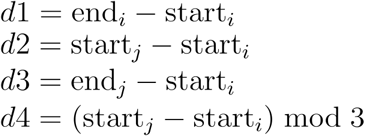

These 4 concatenated distances serve as the input of a small Gated Linear Unit (GLU network), the ‘distance module’, that transforms the 4 dimensional x=[d1, d2, d3, d4] vector into a h dimensional vector (h = Hidden dimension/# attention heads, attention head dimension) that takes the role of 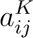 in the relative strand attention.

The distance module takes *x* ∊ R^4^ as input, and applies the following transformations:

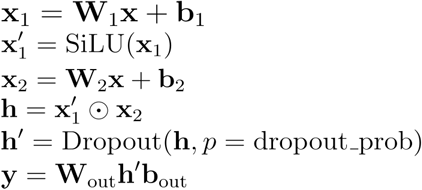

With 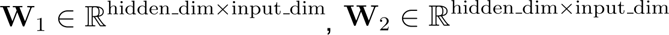 and 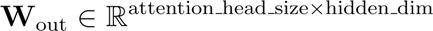. SiLU represents the Sigmoid Linear Unit activation function.

Hyper-parameters choice.

We chose the following hyperparameters for our ALBERT model:

|  |  |
| --- | --- |
| Vocab size | 524,288 |
| Intermediate dimension | 2,048 |
| Embedding dimension | 32 |
| Hidden dimension | 512 |
| # attention heads | 8 |
| # hidden groups | 1 |
| # hidden layers | 6 |
| Distance module GLU intermediate dimension | 32 |
| Distance module dropout prob | 0.1 |

Note that the small embedding size permitted by the ALBERT architecture allows us to use a much larger vocabulary matrix than typically used in BERT models. We also opted to use a single hidden group for the 6 hidden layers, meaning that all encoder layers share weights to reduce the memory footprint and the parameter count. Overall, the ALBERT model is composed of 44M trainable parameters.

### ALBERT-DefenseFinder pretraining

We pre-trained ALBERT on the task of Masked Language Modelling (MLM), adapting the HuggingFace pre-training pipeline.

#### MLM dataset

We sampled N = 6.7 million fragments of up to 64 genes each (432 million tokens) from the Actinomyces genome dataset. To generate a fragment, we first picked a position and a strand at random on one of the contigs, and tried to extend the fragments to the downstream genes, until we either reached the end of the contig or the fragment length reached 64. For each gene in the fragment, we recorded its id, fam50, start, and its coordinates (start, end, strand) on the contig. Furthermore, whenever the gene was annotated as being part of a defense system by DefenseFinder, we recorded the defense system type. Overall, each gene in the corpus was included in 8 fragments on average. 95% of the genomes were used to generate the train set fragments, and the remaining 5% were used for validation.

#### Tokenization

We used a 1 gene = 1 token strategy, using the vocabulary described above to match a gene to a token via its fam50; rare genes, that were not included in the vocabulary, were mapped to the special [UNK] token.

#### Masked Language Modelling

We adapted the AlbertForMaskedLanguageModelling class of HuggingFace to take genomic coordinates as input, and trained our model on the task of masked language modelling using the following settings:

|  |  |
| --- | --- |
| Masking probability | 0.15 |
| Batch size | 32 |
| Gradient accumulation steps | 4 |
| Optimizer | ADAM |
| Learning rate | 5e-4 |
| Learning rate scheduler | Constant with warmup (5000 steps) |
| Mixed precision type | bf16 |
| Number of epochs | 1 |

We monitored the top k accuracy (for k ∈ [1, 5, 10, 100, 1000]) and the cross entropy loss of predictions over the test set fragments every 1000 steps of training. The model with the best loss on the test set was kept for fine-tuning.

### ALBERT-DefenseFinder prediction

#### Fine-tuning datasets

We constructed several datasets for fine-tuning. In the train dataset, we selected all fragments from the MLM dataset containing at least one defense system associated with the “train” defense system types. We first sampled (with replacement) 10,000 fragments from this set, using inverse square root sampling, *i.e.* sampling fragments containing a system s with probability 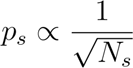, where *N_s_* is the number of times the system was found in the entire dataset, therefore upsampling rare defense systems. We joined these 10,000 fragments with 10,000 fragments uniformly sampled from the MLM dataset. The genes associated with these 20,000 fragments were then assigned a label of 1 if they matched a “train” defense system type, 0 if they didn’t match a defense system, and of −100 if they matched a defense system type from the validation of test sets.

Similarly, we constructed a validation and a test dataset, with 2,000 defensive fragments associated with the validation (respectively test) defense types, sampled as 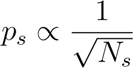 to increase the frequency of rare systems, and 2,000 uniformly sampled fragments. We assigned a label of 1 to genes belonging to a validation (respectively test) defense type, a label of 0 to non defense genes, and a label of −100 to defense genes matched by systems associated to train and test (respectively validation) defense types.

#### Fine-tuning

We adapted the AlbertForTokenClassification class of HuggingFace to take genomic coordinates as input, and trained our model on the task of distinguishing positive from negative labels, at each gene position, ignoring labels −100 over the train dataset fragments. We optimized for a weighted cross entropy loss, weighting examples from each class proportionally to the inverse class frequency. We fine-tuned several models using LoRA finetuning (see section *ESM-DefenseFinder classifier fine-tuning*), and performed hyperparameter optimization using grid-search to maximize the AUROC over the validation dataset over a single epoch. The hyperparameters included:

- Learning rate: [1e-4, 3e-5, 1e-5]
- LoRA rank: [64, 32, 16]
- LoRA factor: [0.5, 1, 2], such that LoRA_α_ = LoRA rank * LoRA factor

After selecting the best hyperparameters (lr=1e-5, LoRA rank = 64, LoRA_α_ = 128), we ran a final fine-tuning using the concatenation of the train and validation test as “train”, and named this model ALBERT-DefenseFinder

#### Inference

We ran ALBERT-DefenseFinder in inference mode to annotate all the fragments in the MLM dataset. We derived a gene token-scope ALBERT-DefenseFinder score using the formula

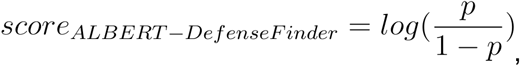

where p is the probability associated by the model to the positive class. These scores were then averaged over fragments to derive a gene-scope ALBERT-DefenseFinder score, and finally these predictions were summarized over fam50 (90th percentile) to derive a family-wise ALBERT-DefenseFinder score that was used to compare the classifier’s prediction to the other methods.

### Construction of multigene systems and selection of candidates

First, we selected the set of all defensive fam50s, and combined them with the set of the 1000 non-defensive families with whose fam50 ALBERT-DefenseFinder scores were the highest (*all_defensive_families*). We computed a graph G with a node per fam50 in this combined set, and a weight per edge reflecting the number of times the two fam50s were seen within a 10-gene window in the Actinomyces selected genomes. We defined the strength of association between two fam50s A and B as:

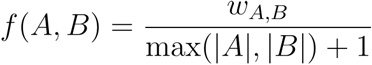

with *w_A,B_* being the weight of the edge between A and B, and |A| and |B| represent the number of times fam A and B were seen in the dataset. We used the Louvain algorithm^58^ to define communities over G, and excluded those that contained known defense systems.

We manually reviewed the list of communities (systems), excluding those that resembled mobile genetic elements (looking for integrases, recombinases or helicases), and focused on systems small enough for DNA synthesis. We manually selected candidates with diverse predicted biochemical functions based on their PFAM annotations.

### Detection of candidates homologs

To identify homologs of the candidate systems identified by Albert-DefenseFinder in Actinomycetes genomes, we first performed a BLAST (version 2.15.0) search starting from the candidate sequences on the RefSeq database^59^. Hits with >80% query coverage and either >40% sequence identity or an E-value <1e-20 were retained and aligned using MAFFT (version v7.520, with --auto settings)^60^. The resulting multiple sequence alignment (MSA) was manually curated, and a Hidden Markov Model (HMM) was built using hmmbuild from the HMMER package, version 3.3.2^51^. We then conducted a second round of sequence search using hmmsearch, applying thresholds of >80% coverage and an E-value <1e-10, followed by alignment with MAFFT. For single-gene systems, the resulting MSA was manually curated and used to construct a new HMM. For multi-gene systems, hits were grouped, aligned with MAFFT, curated, and then used to generate group-specific HMMs. This process was repeated once to refine the detection and alignment of homologous sequences. In the case of Prithvi, where too many hits were found, we determined a GA threshold based on the distribution of scores of 600. Finally, we retrieved taxonomic information for all hits using TaxonKit^61^.

### Strains and growth medium

Bacterial strains and plasmids used in this work are listed in **Supplementary Table 8**. *E. coli* S17 was maintained with LB Miller broth and agar media treated with apramycin (50 μg/mL) when appropriate. *Streptomyces* strains were maintained on Soya-Flour-Mannitol (SFM) agar medium, supplemented with the antibiotics apramycin (50 μg/mL) and Nalidixic Acid (25 μg/mL). Experiments to study phage-bacteria interactions were done in Maltose Yeast Extract Mannitol Medium (4 g/L maltose, 4 g/L yeast extract, 10 g/L malt extract). Difco Nutrient Broth (DNB) medium supplemented with 0.5% glucose and 4 mM Ca(NO3), was used in phage dilutions for the plaque assays.

### Construction of Streptomyces strains for expression of candidate defense systems

The sequences of the selected candidate systems are available in **Supplementary Table 8**. The original coding sequences for each system were refactored into a synthetic operon: all genes from the system were kept in their natural chromosomal organization, a synthetic RBS sequence was placed before the START codon of the first gene, and the operon was put under the control of the constitutive KasOp* promoter. For the system X4 naturally encoded in 2 different convergent operons, the original sequences were kept, and a synthetic RBS and constitutive promoter were placed before the first gene of each operon (KasOp* and ErmE* respectively). The sequences of the designed operons were synthesized by Genscript and cloned into the pOJ436 cosmid vector, between the PvuII and BglII restriction sites. These constructs were then introduced in the chromosome of the strain *S. albus J1074* by conjugation from *Escherichia coli* S17 following standard protocols. Three positive *S. albus* conjugants were screened on SFM medium supplemented with apramycin and the chromosomal integration of the system was confirmed by colony PCR/sequencing.

### Plaque assays

Square plates containing 35 mL of MYM soft agar (0.5% agarose (w/v)) were inoculated with approximately 10⁸ spores of the strain of interest. Plates were dried and 5 μL droplets of serial dilutions of each phage were placed on the agar surface, with dilutions ranging from 10⁻¹ to 10⁻⁷. Plates were incubated at 30°C overnight and placed at room temperature for another 24h, and plaque forming units were counted to evaluate phage replication.

### UMAP

To visualize the space of potential defense systems we selected RefSeq proteins whose representatives at 50% identity and 80% coverage had 10 or more members at 95% identity and 80% coverage. We then selected their associated ESM-DefenseFinder embeddings in 480 dimensions and reduced them to 50 dimensions using a principal component analysis (python module sklearn.decomposition.pca, 84% of the variance explained). We then projected in two dimensions using the python umap^62^ module, and plotted them as a scatter plot with the following color scheme: red for ESM-DefenseFinder scores above the 99th percentile (score 3.27 or more); otherwise dark green for defense scores above the 99th percentile (defense scores 0.286 or more); otherwise light blue for ESM-DefenseFinder scores above the 95th percentile (score 1.13 or more); otherwise light green for defense score above the 95th percentile (defense score 0.106 or more); otherwise gray.

Interactive UMAP is available at https://mdmparis.github.io/antiphage-landscape/. The UMAP is rendered using the scatter-gl library (https://github.com/PAIR-code/scatter-gl). Only the gene family representatives with either defense scores at 30% identity above 0.106, ESM-DefenseFinder scores above 1.13 or which are direct DefenseFinder hits were plotted.

### Protein structure prediction and functional annotation

Protein structures for the candidate defense systems identified in *Streptomyces* and experimentally validated were predicted using AlphaFold3 with default parameters, as available on the AlphaFold Server^42^ (https://golgi.sandbox.google.com/). Protein functional annotation was performed using the HHpred web server^43^ with default parameters. Briefly, a MSA was generated by using an iterative search against UniRef30 using HHblits^43^ (3 MSA generation iterations, minimum probability of 0.20, maximum e-value of 1e-3) and by searching the obtained MSA against the Pfam database^45^ (v37).

### Rarefaction curves

Rarefaction curves were drawn to estimate the capacity of each method (ESM-DefenseFinder and Defense score) to uncover biological novelty among predicted antiphage systems compared to defense systems already mapped by DefenseFinder and PadLoc. To do so, we relied on rarefaction curves, which count how many unique protein families are discovered as additional proteins are predicted as defense systems by each method. When focusing on protein clusters, we filtered out all clusters at 50% identity with less than 5 members and 3 families at 95% identity so that the defense score, which is a statistical measurement, is meaningful. For each predictive method, we considered a protein is predicted as defensive if its assigned score is either greater than (i) the 95th percentile (“stringent” prediction, threshold=0.36 for the defense score method and threshold=3.43 for ESM-DefenseFinder) or than (ii) the 99th percentile (“loose” prediction, threshold=0.13 for the defense score and threshold=1.13 for ESM-DefenseFinder) of the scores distribution. For predicted antiphage proteins, we also filtered out proteins which were predicted as antiphage and already identified by DefenseFinder, to focus on the yet unknown antiphage proteins. The number of protein families was measured using either (i) protein clusters (mmseqs linclust^48^, 50% of identity, 90% coverage, **Figure 4b**) or (ii) the Pfam database of profile HMM^45^ (version 37, **Supplementary Figure 8**). In order to obtain an upper bound for the rate of discovery of protein families as new proteins are predicted as antiphage, we also applied the same approach by randomly sampling proteins from the RefSeq database of prokaryotic complete genomes. Each rarefaction curve was fitted using a Michaelis-Menten model with an intercept 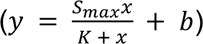 to estimate (i) its asymptote *S_max_* and (ii) its initial discovery rate of protein families *V*_0_ = *S_max_* / *K* which corresponds to the slope of the curve when the number of predicted antiphage proteins *x* tends to 0. The curve was fitted by minimizing the Root Mean-Squared Deviation (RMSD) between the empirical and the fitted curves using the minimize function from the scipy^63^ python package (version 1.14.1). For each method, the asymptote Smax estimates the total number of protein families involved in antiphage defense (**Figure 4c**) and the initial discovery rate V0 estimates the initial novelty yielded by the approach compared to what is already known in DefenseFinder.

## Acknowledgements

We are grateful to Vincent Libis (INSERM U1284) for the gift of the strain *S. albus J1074* and the pOJ436 vector. Several bioinformatic analyses were performed on the Core Cluster of the Institut Français de Bioinformatique (IFB) (ANR-11-INBS-0013).

We would like to thank Vincent Libis, Ariel Lindner, Enzo Poirier, Lucas Paoli, Francois Rousset, Adi Millman for fruitful discussions and feedback.

## Authors contributions

E.M. and A.B conceptualized the project. E.M., A.H., H.V., F.T. and J. C. carried out all the bioinformatic analyses. E.M. and H.S. chose and designed strains; T.C., R.L. performed experiments to assess anti-phage activity. A.B. supervised the project. E.M., and A.B. wrote the initial version of the manuscript. All authors contributed to the design of the experiments, the discussion of the results and the final version of the manuscript.

## Data availability

Supplementary materials, data and code are available at https://github.com/mdmparis/antiphage_landscape_2025

## References

1. Georjon, H. & Bernheim, A. The highly diverse antiphage defence systems of bacteria. Nat. Rev. Microbiol. 21, 686–700 (2023).

2. Tesson, F. et al. A Comprehensive Resource for Exploring Antiphage Defense: DefenseFinder Webservice,Wiki and Databases. Peer Community J. 4, (2024).

3. Hu, H. et al. Structure and mechanism of the Zorya anti-phage defense system. Nature (2024) doi:10.1038/s41586-024-08493-8.

4. Cohen, D. et al. Cyclic GMP-AMP signalling protects bacteria against viral infection. Nature 574, 691–695 (2019).

5. Ofir, G. et al. Antiviral activity of bacterial TIR domains via immune signalling molecules. Nature 600, 116–120 (2021).

6. Wilkinson, M. E., Li, D., Gao, A., Macrae, R. K. & Zhang, F. Phage-triggered reverse transcription assembles a toxic repetitive gene from a noncoding RNA. Science 386, eadq3977 (2024).

7. Johnson, A. G. et al. Bacterial gasdermins reveal an ancient mechanism of cell death. Science 375, 221–225 (2022).

8. Tang, S. et al. De novo gene synthesis by an antiviral reverse transcriptase. Science 386, eadq0876 (2024).

9. Makarova, K. S., Wolf, Y. I., Snir, S. & Koonin, E. V. Defense Islands in Bacterial and Archaeal Genomes and Prediction of Novel Defense Systems. J. Bacteriol. 193, 6039–6056 (2011).

10. Doron, S. et al. Systematic discovery of antiphage defense systems in the microbial pangenome. Science 359, eaar4120 (2018).

11. Gao, L. et al. Diverse enzymatic activities mediate antiviral immunity in prokaryotes. Science 369, 1077–1084 (2020).

12. Millman, A. et al. An expanded arsenal of immune systems that protect bacteria from phages. Cell Host Microbe 30, 1556–1569.e5 (2022).

13. Rousset, F. et al. Phages and their satellites encode hotspots of antiviral systems. Cell Host Microbe 30, 740–753.e5 (2022).

14. Fillol-Salom, A. et al. Bacteriophages benefit from mobilizing pathogenicity islands encoding immune systems against competitors. Cell 185, 3248–3262.e20 (2022).

15. Darracq, B. et al. Sedentary chromosomal integrons as biobanks of bacterial anti-phage defence systems. 2024.07.02.601686 Preprint at 10.1101/2024.07.02.601686 (2024).

16. Kieffer, N. et al. Mobile Integrons Encode Phage Defense Systems. Preprint at 10.1101/2024.07.02.601719 (2024).

17. Payne, L. J., Hughes, T. C. D., Fineran, P. C. & Jackson, S. A. New antiviral defences are genetically embedded within prokaryotic immune systems. 2024.01.29.577857 Preprint at 10.1101/2024.01.29.577857 (2024).

18. Zaworski, J. et al. Reassembling a cannon in the DNA defense arsenal: Genetics of StySA, a BREX phage exclusion system in Salmonella lab strains. PLoS Genet. 18, e1009943 (2022).

19. Vassallo, C. N., Doering, C. R., Littlehale, M. L., Teodoro, G. I. C. & Laub, M. T. A functional selection reveals previously undetected anti-phage defence systems in the E. coli pangenome. Nat. Microbiol. 7, 1568–1579 (2022).

20. Georjon, H., Tesson, F., Shomar, H. & Bernheim, A. Genomic characterization of the antiviral arsenal of Actinobacteria: This article is part of the Microbial Evolution collection. Microbiology 169, (2023).

21. Shomar, H. et al. A family of lanthipeptides with anti-phage function. bioRxiv 2024–06 (2024).

22. Naveed, H., et al. A Comprehensive Overview of Large Language Models. Preprint at 10.48550/arXiv.2307.06435 (2024).

23. Vaswani, A., et al. Attention Is All You Need. Preprint at 10.48550/arXiv.1706.03762 (2023).

24. Rives, A. et al. Biological structure and function emerge from scaling unsupervised learning to 250 million protein sequences. Proc. Natl. Acad. Sci. U. S. A. 118, e2016239118 (2021).

25. Cornman, A. et al. The OMG dataset: An Open MetaGenomic corpus for mixed-modality genomic language modeling. 2024.08.14.607850 Preprint at 10.1101/2024.08.14.607850 (2024).

26. Hwang, Y., Cornman, A. L., Kellogg, E. H., Ovchinnikov, S. & Girguis, P. R. Genomic language model predicts protein co-regulation and function. Nat. Commun. 15, 2880 (2024).

27. Miller, D., Stern, A. & Burstein, D. Deciphering microbial gene function using natural language processing. Nat. Commun. 13, 5731 (2022).

28. Tesson, F. et al. Systematic and quantitative view of the antiviral arsenal of prokaryotes. Nat. Commun. 13, (2022).

29. Payne, L. J. et al. Identification and classification of antiviral defence systems in bacteria and archaea with PADLOC reveals new system types. Nucleic Acids Res. 49, 10868–10878 (2021).

30. Kim, G. B. et al. Functional annotation of enzyme-encoding genes using deep learning with transformer layers. Nat. Commun. 14, 7370 (2023).

31. Garb, J. et al. Multiple phage resistance systems inhibit infection via SIR2-dependent NAD+ depletion. Nat. Microbiol. 7, 1849–1856 (2022).

32. Bernheim, A. et al. Inhibition of NHEJ repair by type II-A CRISPR-Cas systems in bacteria. Nat. Commun. 8, 2094 (2017).

33. Ramisetty, B. C. M., Natarajan, B. & Santhosh, R. S. mazEF-mediated programmed cell death in bacteria: ‘what is this?’ *Crit*. Rev. Microbiol. 41, 89–100 (2015).

34. Kumar, R. et al. The bacterial condensin MukB compacts DNA by sequestering supercoils and stabilizing topologically isolated loops. J. Biol. Chem. 292, 16904–16920 (2017).

35. Broadbent, S. E., Balbontin, R., Casadesus, J., Marinus, M. G. & van der Woude, M. YhdJ, a nonessential CcrM-like DNA methyltransferase of Escherichia coli and Salmonella enterica. J. Bacteriol. 189, 4325–4327 (2007).

36. Dot, E. W., Thomason, L. C. & Chappie, J. S. Everything OLD is new again: How structural, functional, and bioinformatic advances have redefined a neglected nuclease family. Mol. Microbiol. 120, 122–140 (2023).

37. Burman, N. et al. A virally encoded tRNA neutralizes the PARIS antiviral defence system. Nature 634, 424–431 (2024).

38. Sundararajan, M., Taly, A. & Yan, Q. Axiomatic Attribution for Deep Networks. Preprint at 10.48550/arXiv.1703.01365 (2017).

39. Lan, Z., et al. ALBERT: A Lite BERT for Self-supervised Learning of Language Representations. Preprint at 10.48550/arXiv.1909.11942 (2020).

40. Devlin, J., Chang, M.-W., Lee, K. & Toutanova, K. BERT: Pre-training of Deep Bidirectional Transformers for Language Understanding. Preprint at 10.48550/arXiv.1810.04805 (2019).

41. Aframian, N., Omer Bendori, S., Hen, T., Guler, P. & Eldar, A. High defense system expression broadens protection range at the cost of increased autoimmunity. Preprint at 10.1101/2023.11.30.569366 (2023).

42. Abramson, J. et al. Accurate structure prediction of biomolecular interactions with AlphaFold 3. Nature 630, 493–500 (2024).

43. Zimmermann, L. et al. A Completely Reimplemented MPI Bioinformatics Toolkit with a New HHpred Server at its Core. J. Mol. Biol. 430, 2237–2243 (2018).

44. Gabler, F., et al. Protein Sequence Analysis Using the MPI Bioinformatics Toolkit. Curr. Protoc. Bioinforma. 72, e108 (2020).

45. Mistry, J. et al. Pfam: The protein families database in 2021. Nucleic Acids Res. 49, D412–D419 (2021).

46. Kibby, E. M. et al. Bacterial NLR-related proteins protect against phage. Cell 186, 2410–2424.e18 (2023).

47. O’Leary, N. A. et al. Reference sequence (RefSeq) database at NCBI: current status, taxonomic expansion, and functional annotation. Nucleic Acids Res. 44, D733–745 (2016).

48. Hauser, M., Steinegger, M. & Söding, J. MMseqs software suite for fast and deep clustering and searching of large protein sequence sets. Bioinforma. Oxf. Engl. 32, 1323–1330 (2016).

49. Ondov, B. D. et al. Mash: fast genome and metagenome distance estimation using MinHash. Genome Biol. 17, 132 (2016).

50. Seemann, T. Prokka: rapid prokaryotic genome annotation. Bioinforma. Oxf. Engl. 30, 2068–2069 (2014).

51. Eddy, S. R. Accelerated Profile HMM Searches. PLOS Comput. Biol. 7, e1002195 (2011).

52. Hu, E. J., et al. LoRA: Low-Rank Adaptation of Large Language Models. Preprint at 10.48550/arXiv.2106.09685 (2021).

53. Schmirler, R., Heinzinger, M. & Rost, B. Fine-tuning protein language models boosts predictions across diverse tasks. Nat. Commun. 15, 7407 (2024).

54. Hayes, T. et al. Simulating 500 million years of evolution with a language model. 2024.07.01.600583 Preprint at 10.1101/2024.07.01.600583 (2024).

55. van Kempen, M. et al. Fast and accurate protein structure search with Foldseek. Nat. Biotechnol. 42, 243–246 (2024).

56. Mikolov, T., Sutskever, I., Chen, K., Corrado, G. & Dean, J. Distributed Representations of Words and Phrases and their Compositionality. Preprint at 10.48550/arXiv.1310.4546 (2013).

57. Shaw, P., Uszkoreit, J. & Vaswani, A. Self-Attention with Relative Position Representations. Preprint at 10.48550/arXiv.1803.02155 (2018).

58. Blondel, V. D., Guillaume, J.-L., Lambiotte, R. & Lefebvre, E. Fast unfolding of communities in large networks. J. Stat. Mech. Theory Exp. 2008, P10008 (2008).

59. Camacho, C. et al. BLAST+: architecture and applications. BMC Bioinformatics 10, 421 (2009).

60. Katoh, K., Rozewicki, J. & Yamada, K. D. MAFFT online service: multiple sequence alignment, interactive sequence choice and visualization. Brief. Bioinform. 20, 1160–1166 (2019).

61. Shen W. & Ren H. TaxonKit: A practical and efficient NCBI taxonomy toolkit. J. Genet. Genomics 48, 844–850 (2021).

62. Sainburg, T., McInnes, L. & Gentner, T. Q. Parametric UMAP embeddings for representation and semi-supervised learning. Preprint at 10.48550/arXiv.2009.12981 (2021).

63. Virtanen, P. et al. SciPy 1.0: fundamental algorithms for scientific computing in Python. Nat. Methods 17, 261–272 (2020).

