## Supplementary figures for "Protein and genomic language models chart a vast landscape of antiphage defenses"

### Supplementary Materials Mordret et al

#### List of supplementary Tables

**Supplementary Table 1: DefenseFinder results on RefSeq.** Number of each system found by DefenseFinder in the 32,798 genomes of the RefSeq complete genome database.

**Supplementary Table 2: PadLoc results on RefSeq.** Number of each system found by PadLoc in the 32,800 genomes of the RefSeq complete genome database.

**Supplementary Table 3: Defense system type split for ESM training and evaluation datasets.** Association between each of the 137 defense types and Train/Eval/Test

**Supplementary Table 4 ESM and DefenseScore results on the RefSeq database.** Results of ESM and DefenseScore for 50% identity (80% coverage) with ESM score >= 1.077 or DefenseScore >= 0.2 and clusters with more than 5 members and 3 different representatives at 95% identity (n = 175,756) .

**Supplementary Table 5 ESM results of the pangenome of Escherichia coli:** Results of ESM on all *E. coli* representative proteins clustered at 80% identity.

**Supplementary Table 6 Results of the different models on all Actinomycetota protein clusters at 50% identity.**

**Supplementary Tables 7 List of all candidates defense systems with the different scores.**
All sequences of experimentally tested and validated defense systems with their scores with the different approaches.

**Supplementary Tables 8 List of all strains, vectors and phage used in the study**

#### Supplementary Figures

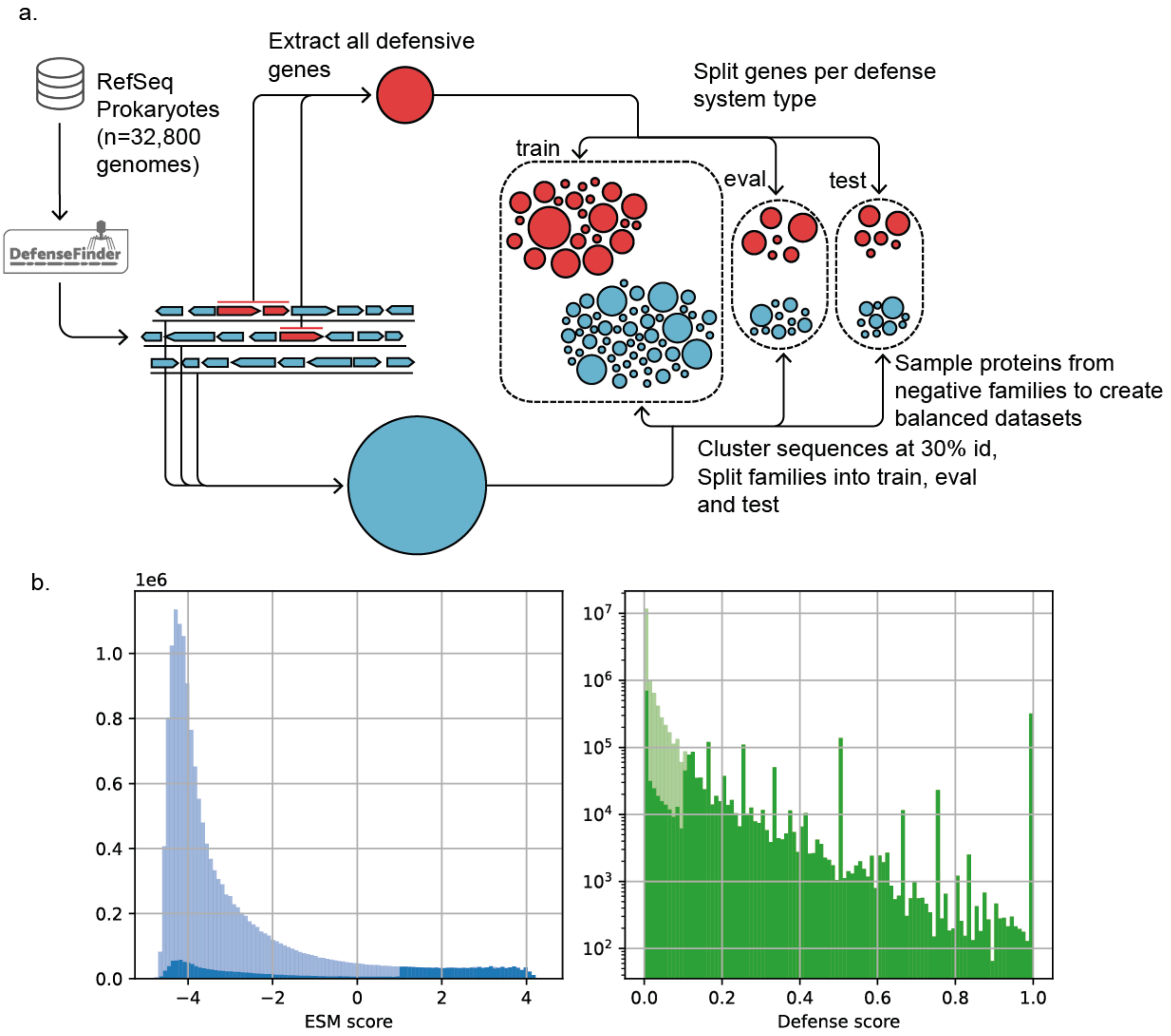


**Supplementary Figure 1: ESM-DefenseFinder training**

a. ESM training workflow for ESM-DefenseFinder b. Distribution of ESM-DefenseFinder (left) and Defense Score (right). Light colors: distribution of scores over all RefSeq. Dark colors: distribution of scores over high scoring proteins (ESM score > 1.077 OR Defense score > 0.106).

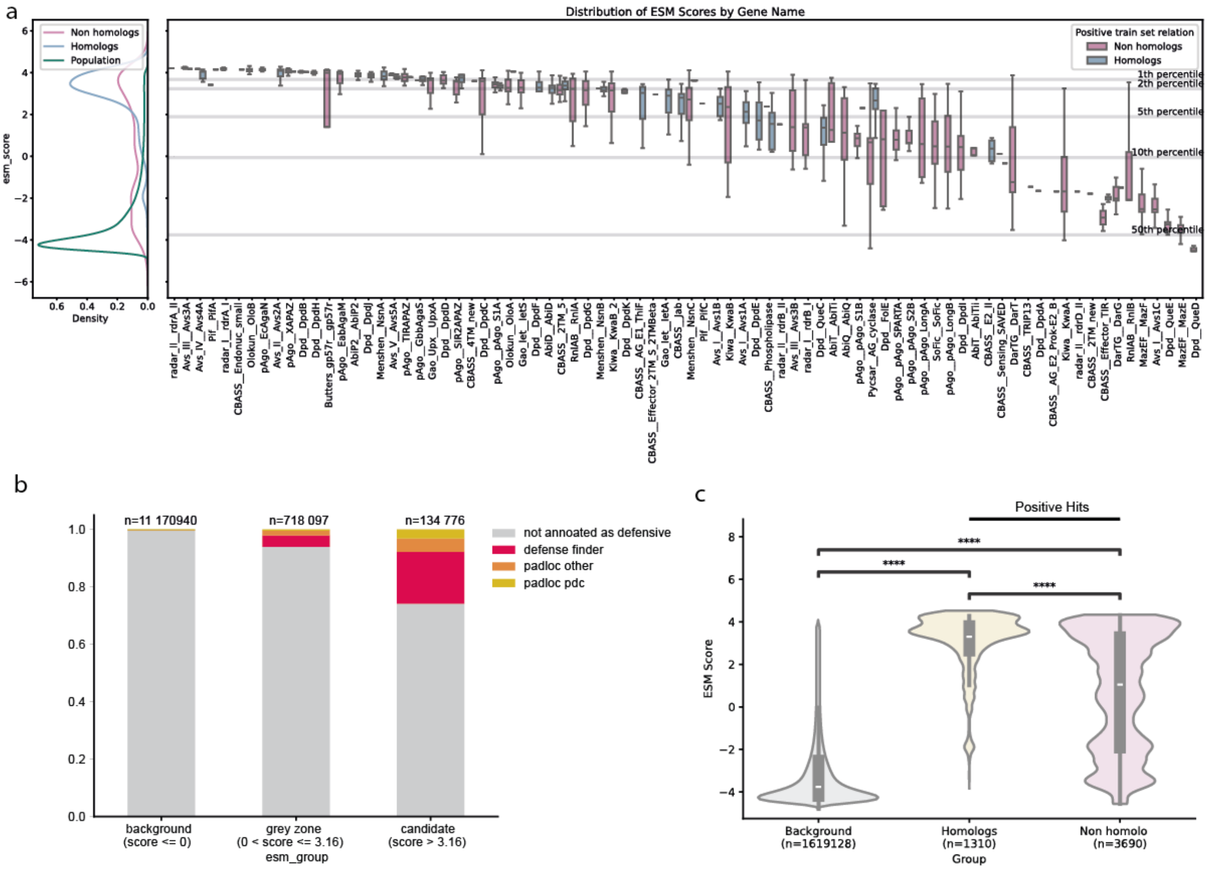


**Supplementary Figure 2: ESM-DefenseFinder relies on homology**

**a.** Model's response over genes of the test set grouped by the HMM profile that matched them (x-axis) and whether they showed signatures of homology to at least one of the positive train set proteins (blue: detected homology, pink: no sign of homology). Vertical lines represent the percentile of background distribution **b.** High scoring hits are enriched for defense systems. The 12M genes from *E. coli* genomes in the RefSeq dataset were grouped by ESM-DefenseFinder score (as computed on their representative fam80) into 3 bins, representing low scoring (ESM-DefenseFinder score < 0), medium scoring (0 <= ESM-DefenseFinder score < 3.16) or high scoring (ESM-DefenseFinder score > 3.16) genes. For each bin, we recorded the proportion of defensive and non defensive proteins, annotating first with DefenseFinder (pink), and then annotating the remaining genes with PadLoc. PadLoc annotations were split between PadLoc PDC (candidate systems, not experimentally verified, yellow) and PadLoc others (orange) **c.** Distribution of ESM-DefenseFinder scores for the negative ("Background") and positives families from the test set, at 80% aa identity. These positive families were further split based on whether or not they showed marks of homology to positive train set families ("Homologs" and "Non Homologs"). Homology was assessed via a sequence search on the predicted structures of the train and test set positive proteins, using MMSEQs-search. The two-sided Mann-Whitney U test was used to compare the distributions of the populations, with *** indicating a p-value < 0.001.

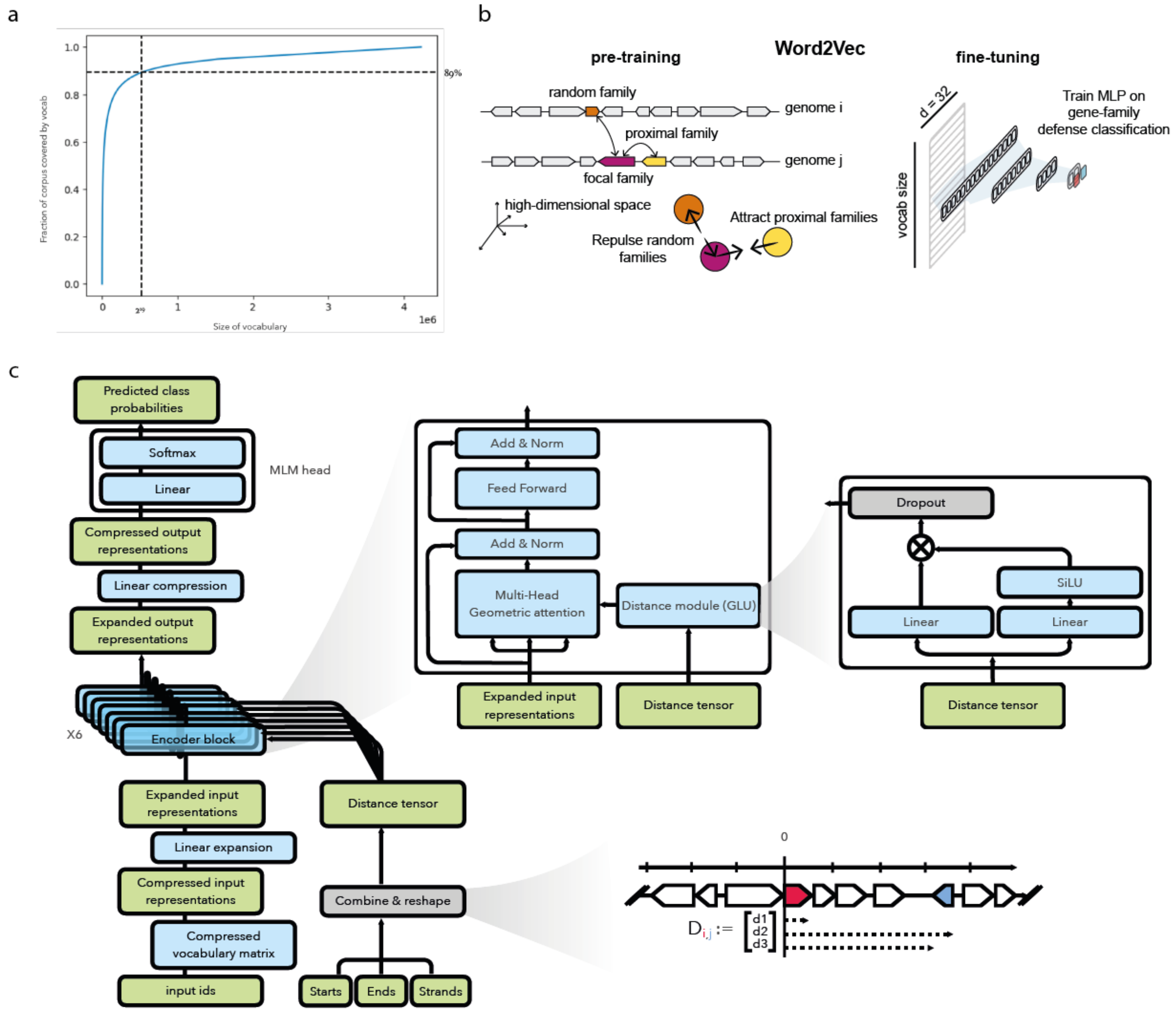


**Supplementary Figure 3: Context based methods workflow**

**a. Vocabulary.** Fraction of the corpus covered at various vocabulary size thresholds. We ranked genes by fam50 size, and estimated what fraction of the total corpus (y-axis) would be included if using only the top N biggest fam50 (x-axis). At the chosen vocabulary size ($2^{19}$ families), we include 89% of the genes. **b. Schematic representation of the Word2Vec training.** Left: pretraining. Each entry in the vocabulary is initialized with a random vector. During the training procedure, pairs of families are chosen, either as “context pairs” (focal family, purple, and proximal family , yellow) or “random pairs” (focal family + random family, orange). The optimization procedure aims at bringing closer context pairs (low cosine distance) while replusing “random pairs” (high cosine distance). Right: transfer learning. We use the frozen embeddings of Word2Vec as input to Multi Layer Perceptrons, that we optimize on fam50 defense classification using cross entropy loss. **c. ALBERT.** Left: model architecture. The architecture is a variant of ALBERT, an encoder-only transformer that stores its vocabulary matrix in a compressed format ($d_e$=embedding dimension), before expanding it to the hidden dimension $d_h$. The data then flows through 6 encoder blocks whose weights are tied, before being compressed from $d_h$ to $d_e$. During MLM, the token representations in dimension $d_e$ are then projected onto vocabulary, minimizing cross entropy loss at masked positions. In addition to the genes' input ids, the encoder stack also takes as input genomic coordinates for each gene in the fragment. Global starts, ends strands of genes are first combined and reshaped to form a *distance tensor* of shape (batch size, sequence length, sequence length, 4), where the distances between the boundaries of genes i and j are stored as *distance tensor* [b,i,j,n] = $d_{n,i,j}$. Distances measure the difference in coordinates between the start of $gene_i$, and either the end of $gene_i$ ($d_1$), the start of $gene_j$ ($d_2$) or the end of $gene_j$ ($d_3$), in the local reference frame defined by the start and the strand of $gene_i$. A 4th distance, $d_4$, captures whether the genes are in frame ($d_4 = d_2$ % 3). The *distance tensor* is projected in dimension attention head dimension $d_{ah}$ by a Gated Liner Unit (GLU) that we call the *distance module*, and the resulting tensor serves as input to the relative key attention of the encoder stack.

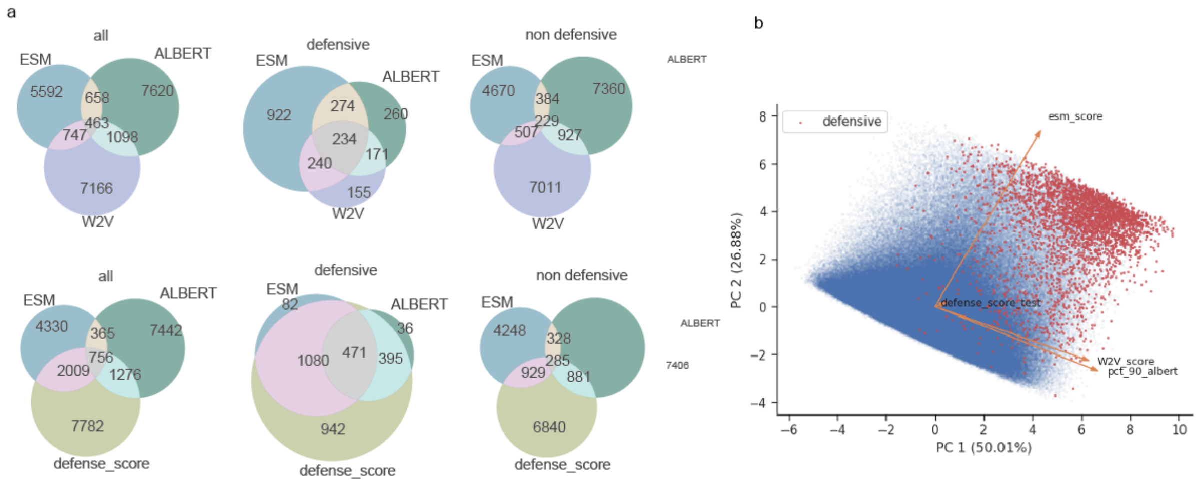


**Supplementary Figure 4: Comparison of prediction methods**: **a.** Venn diagrams of the test set fam50s predicted as defensive by the different methods, using their respective best F1-score as cutoffs, either will all genes (left), or splitting the data between fam50s known to be defensive (middle) or not (right). **b** PCA of the 4 scores on all genes in the dataset of actinobacteria.

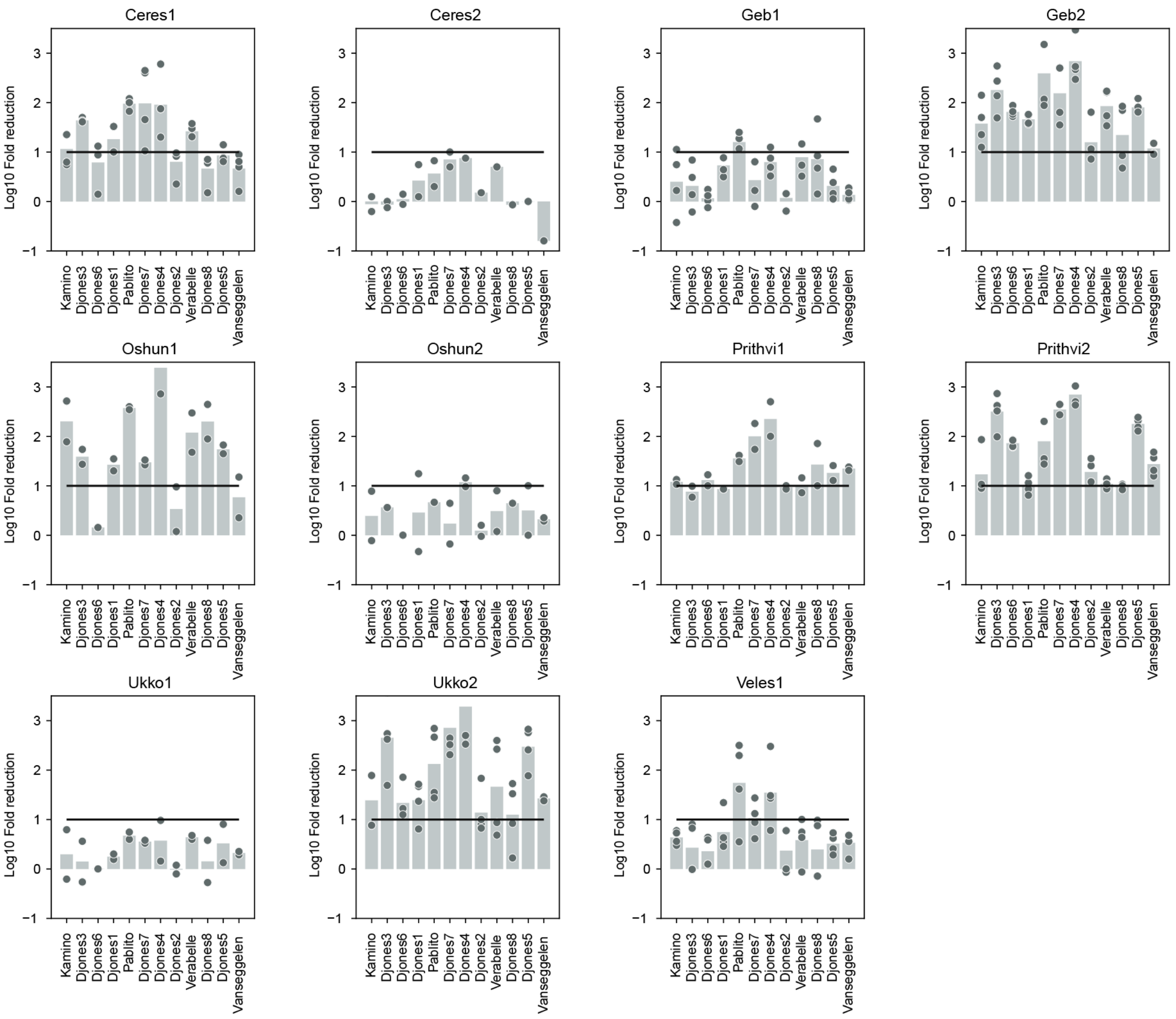


**Supplementary Figure 5: Experimental validation of Streptomyces candidates systems**
Log10 fold reduction of the efficiency of plating of phages infecting candidate strains. Negative control was constructed in the same way but expressing enzymes lacking antiphage activity (Methods). Bar graphs represent an average of four biological replicates, with individual data points overlaid. An horizontal line is represented for Log10 fold of 1, which was used to determine the antiphage activity.

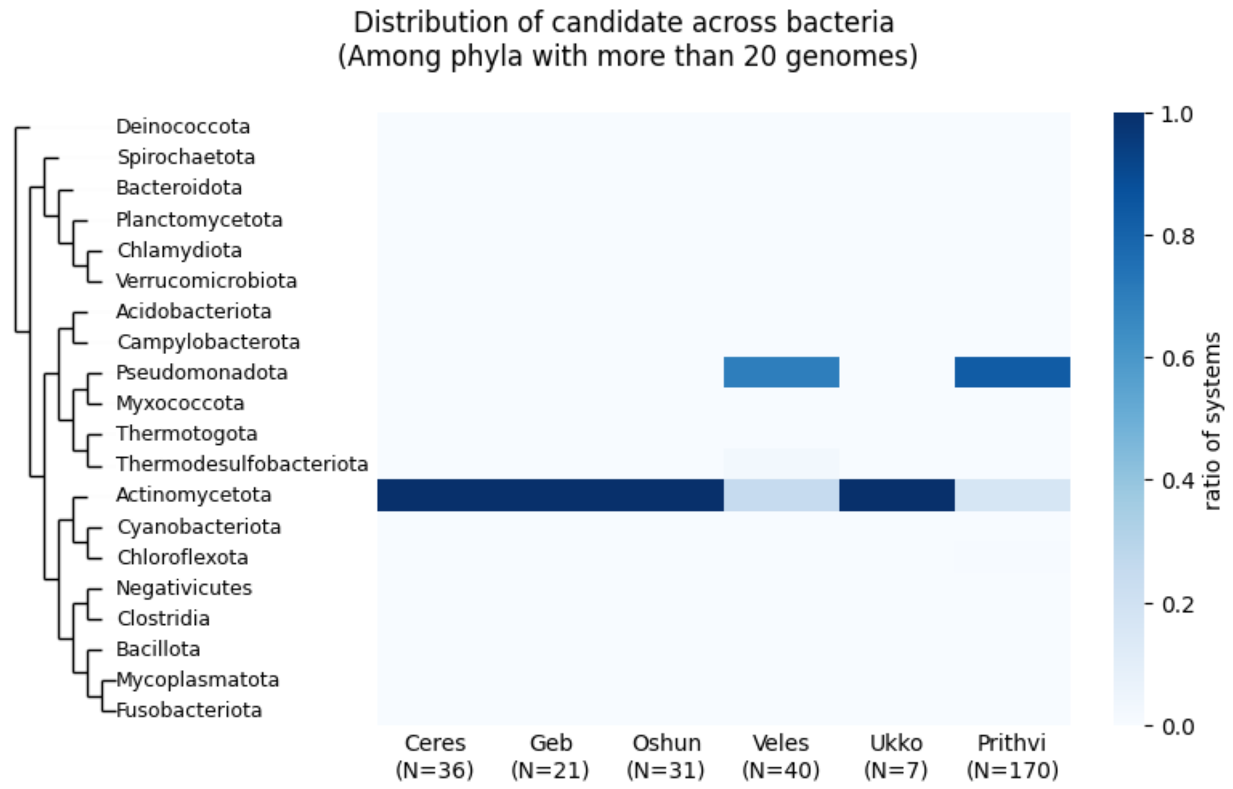


**Supplementary Figure 6: Taxonomic Distribution of novel Streptomyces antiphage systems**

For each system, number of homologs is indicated. Ratio of systems means number of systems in clade out of total number of homologs.

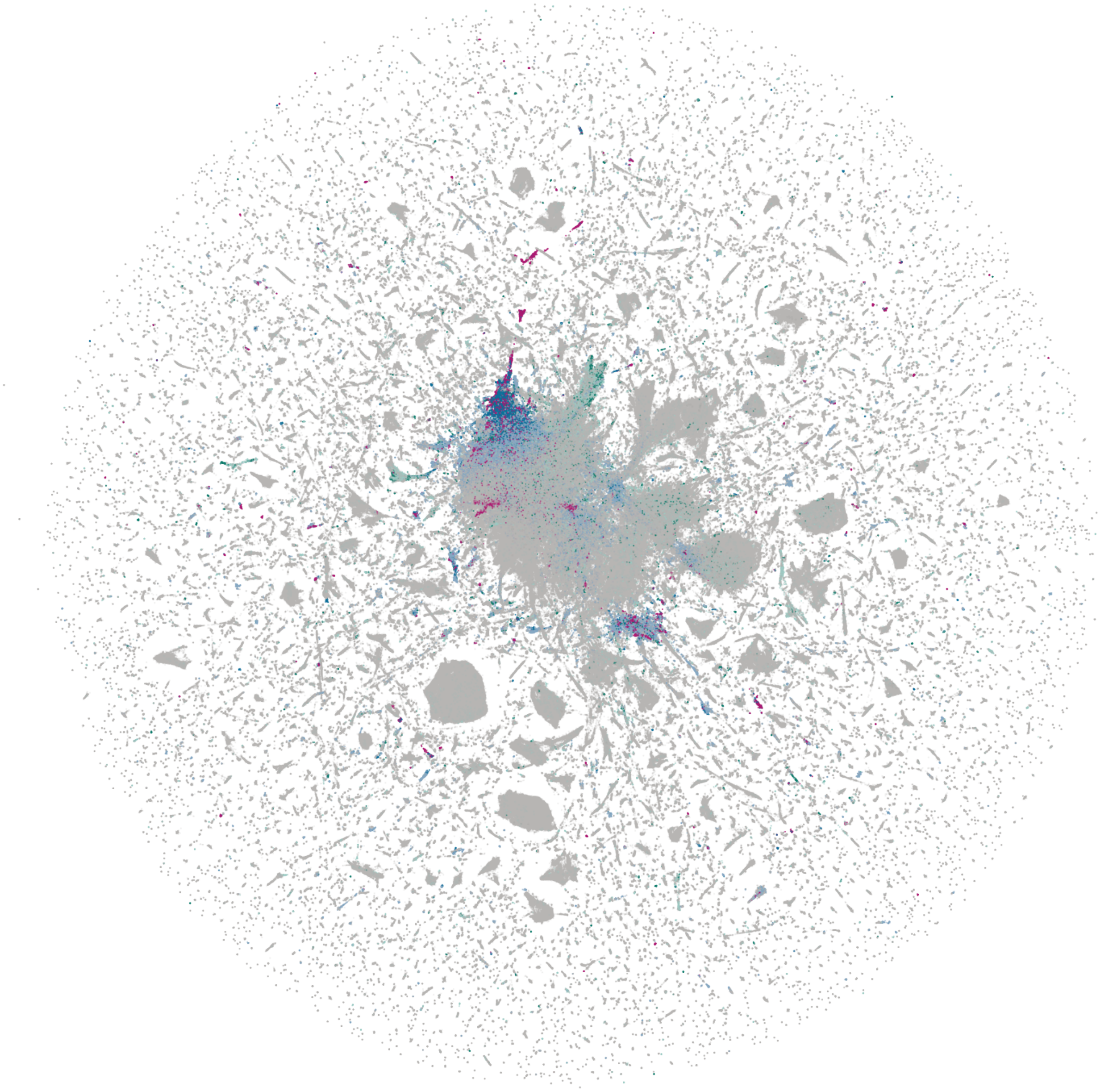

**Supplementary Figure 7: Full UMAP**

Projection of ESM-DefenseFinder embeddings of families clustered at 50% identity, 50% coverage with more than 10 members. Pink: families with a member annotated by DefenseFinder. Blue: families predicted as antiphage by ESM-DefenseFinder (dark blue, score > best F1-score, light blue score >95th percentile). Green: families predicted as antiphage by Defense Score (dark green, score > best F1-score, light green score >95th percentile).

An interactive version is available at [https://mdmparis.github.io/antiphage-landscape/](https://mdmparis.github.io/antiphage-landscape/)

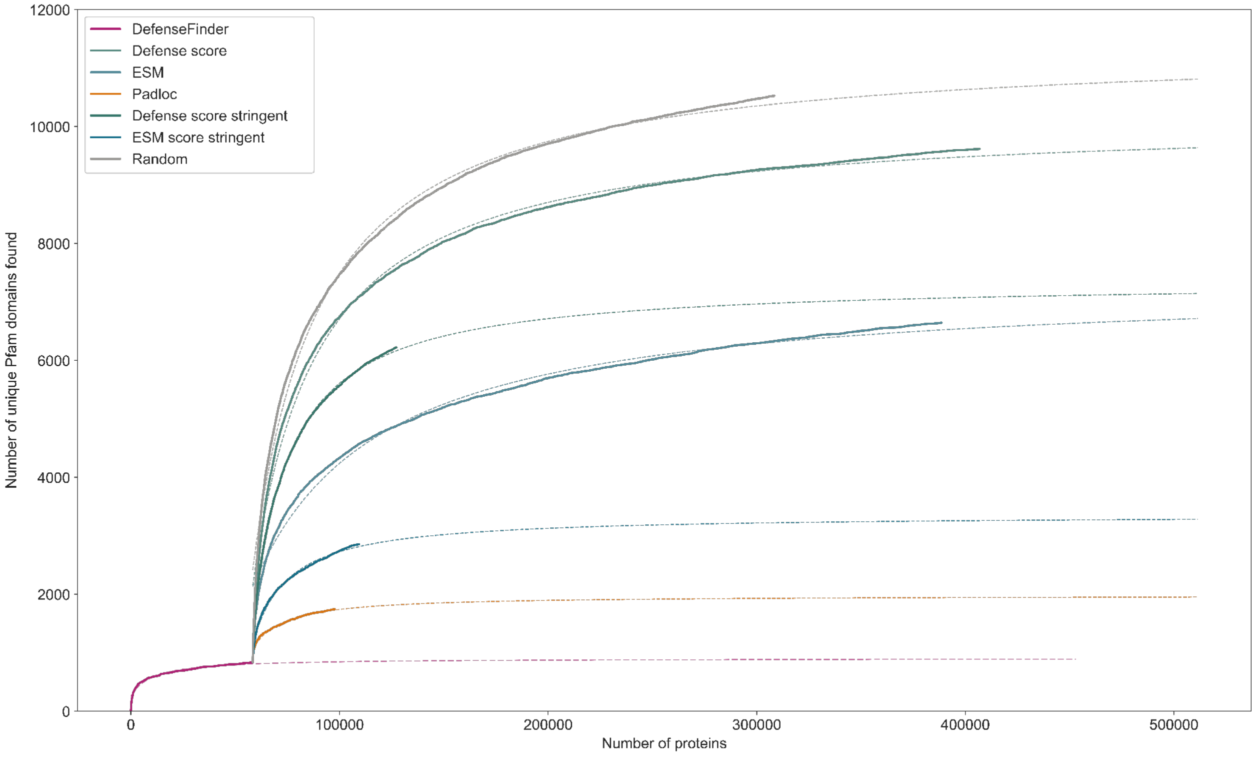


**Supplementary Figure 8: Rarefaction curves of PFAMs in predicted antiphage protein families**

Rarefaction curves measuring how many novel protein domains from Pfam are discovered as new proteins are detected as antiphage according to DefenseFinder (pink) and PadLoc (orange), or predicted as antiphage according to the ESM score (stringent prediction: dark blue, loose prediction: light blue) or the Defense score (stringent prediction: dark green, loose prediction: light green). For each method, the best fit using a Michaelis-Menten model is drawn in dashed lines.
